# A human RNA ligase that operates via auto- and RNA-AMPylation

**DOI:** 10.1101/2022.07.18.500566

**Authors:** Yizhi Yuan, Florian M. Stumpf, Lisa A. Schlor, Olivia P. Schmidt, Luisa B. Huber, Matthias Frese, Eva Höllmüller, Martin Scheffner, Florian Stengel, Kay Diederichs, Andreas Marx

## Abstract

Different forms of life are known to express RNA ligases that catalyse the condensation of a 3’-hydroxy group and a 5’-terminal phosphate of RNA. No such RNA ligases have yet been identified in vertebrates. Here, we report that the hitherto uncharacterised human protein chromosome 12 open reading frame 29 (C12orf29), which we identified by a chemical proteomics approach, is a 5’-3’ RNA ligase. C12orf29 catalyses RNA ligation via auto-AMPylation of a critical lysine residue by using ATP as a cosubstrate and subsequent AMP transfer to the 5’-phosphate of an RNA substrate followed by phosphodiester bond formation. Studies at the cellular level reveal the involvement of C12orf29 in maintaining RNA integrity upon cellular stress induced by reactive oxygen species. These findings highlight the importance of RNA ligation for cellular fitness.

## Main Text

DNA ligases seal nicks between a 3’-hydroxy group and a 5’-terminal phosphate in DNA ^1-3^. They are indispensable for maintaining genome integrity due to their involvement in DNA replication, recombination and repair throughout all domains of life. In several forms of life, but not in higher eukaryotes such as vertebrates, RNA ligases that catalyse the corresponding reaction with RNA have been described and linked to RNA repair^4-6^. Surprisingly, proteins with 5’-3’ RNA ligase activity have not yet been identified in higher eukaryotes, such as humans, although cell culture experiments conducted already in 1983 suggested their presence^7^. Here, we report the first human enzyme of that kind by showing that the hitherto uncharacterised human protein chromosome 12 open reading frame 29 (C12orf29) is a 5’-3’ RNA ligase.

### Discovery of C12orf29 by chemical proteomics dedicated to identify AMPylated proteins

AMPylation is a posttranslational modification (PTM) in which AMP is covalently attached to an amino acid side chain of target proteins via a phosphodiester bond using ATP as cosubstrate^8,9^. In order to gain further insights into the physiological roles of AMPylation, we devised a chemical proteomics approach (Fig. 1a-c) to discover AMPylation targets by using a chemical probe based on diadenosine triphosphate (Ap_3_A), a naturally occurring nucleotide that is formed increasingly in response to cellular stress^10^. From the proteins identified by affinity purification mass spectrometry (AP-MS), one protein caught our attention since it had not yet been characterized: C12orf29. We next verified this protein by immunoblotting after affinity purification from human embryonic kidney cells (HEK293T) and human non-small cell lung carcinoma cells (H1299) (Fig. 1d).

**Fig. 1.**
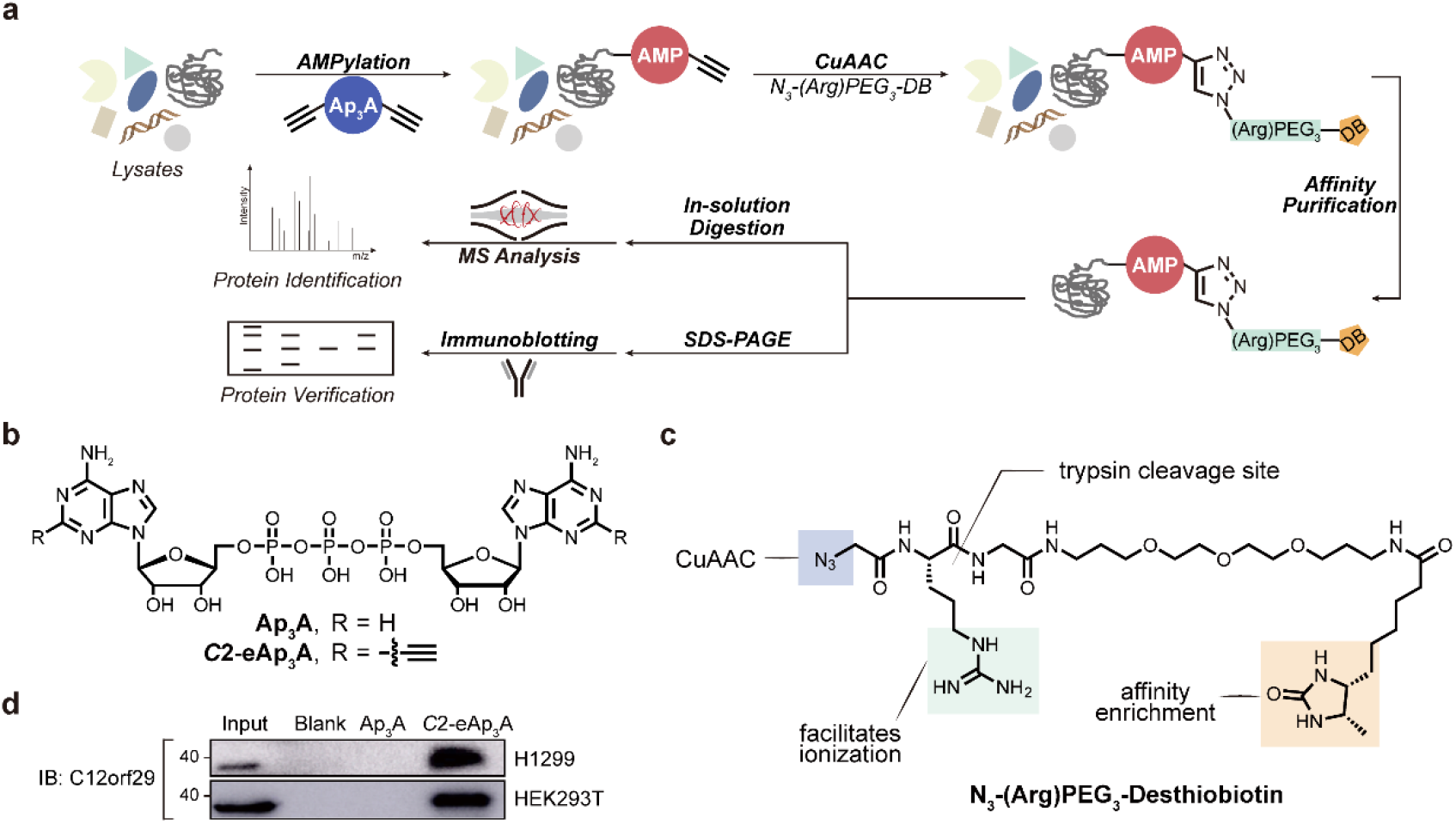
Identification of C12orf29 by chemical proteomics using modified nucleotide probes. **a**, Schematic display of the workflow for the identification of C12orf29. Cell lysates were incubated with *C*2-eAp_3_A (structure see Fig. 1b) or controls. AMPylated proteins are expected to bear ethynyl functionalities that enabled selective modification with an affinity tag desthiobiotin (DB) via copper(I)-catalysed azide alkyne cycloaddition (CuAAC). Labelled proteins were purified and identified by AP-MS, and further verified by immunoblotting. **b**, Structures of Ap_3_A analogues employed in this study. **c**, Structure of the azide bearing desthiobiotin as affinity tag. **d**, Affinity purification of C12orf29 from two cell lysates verified by immunoblotting.

### C12orf29 is an RNA ligase operating via sequential auto- and RNA AMPylation

C12orf29 is a 37 kDa human protein consisting of 325 amino acids. Sequence analysis shows that it is highly conserved among higher eukaryotes like vertebrates (Supplementary Fig. 1) but absent in lower eukaryotes species such as yeast. To obtain insight into its potential function, we initially performed structure prediction of C12orf29 using bioinformatic tools. Intriguingly, a putative structure of a truncated form of C12orf29 was obtained by Phyre2^11^ based on the identity of 22 out of 71 residues (195-265) with those of *Naegleria gruberi* RNA ligase (*Ngr*Rnl)^12,13^. *Ngr*Rnl is a 5’-3’ RNA ligase and as such operates via a three-step mechanism alike DNA ligases (Fig. 2a)^4,6^. First, the ligase auto-AMPylates the catalytic lysine residue by the use of ATP resulting in the ligase-(lysyl-N)-AMP species. Then, AMP is transferred onto the 5’-phosphate end of RNA (pRNA) to yield the RNA-adenylate intermediate (AppRNA). Finally, the two RNA ends are ligated by phosphodiester bond formation upon nucleophilic attack of the 3’-OH to the AppRNA to liberate AMP.

**Fig. 2.**
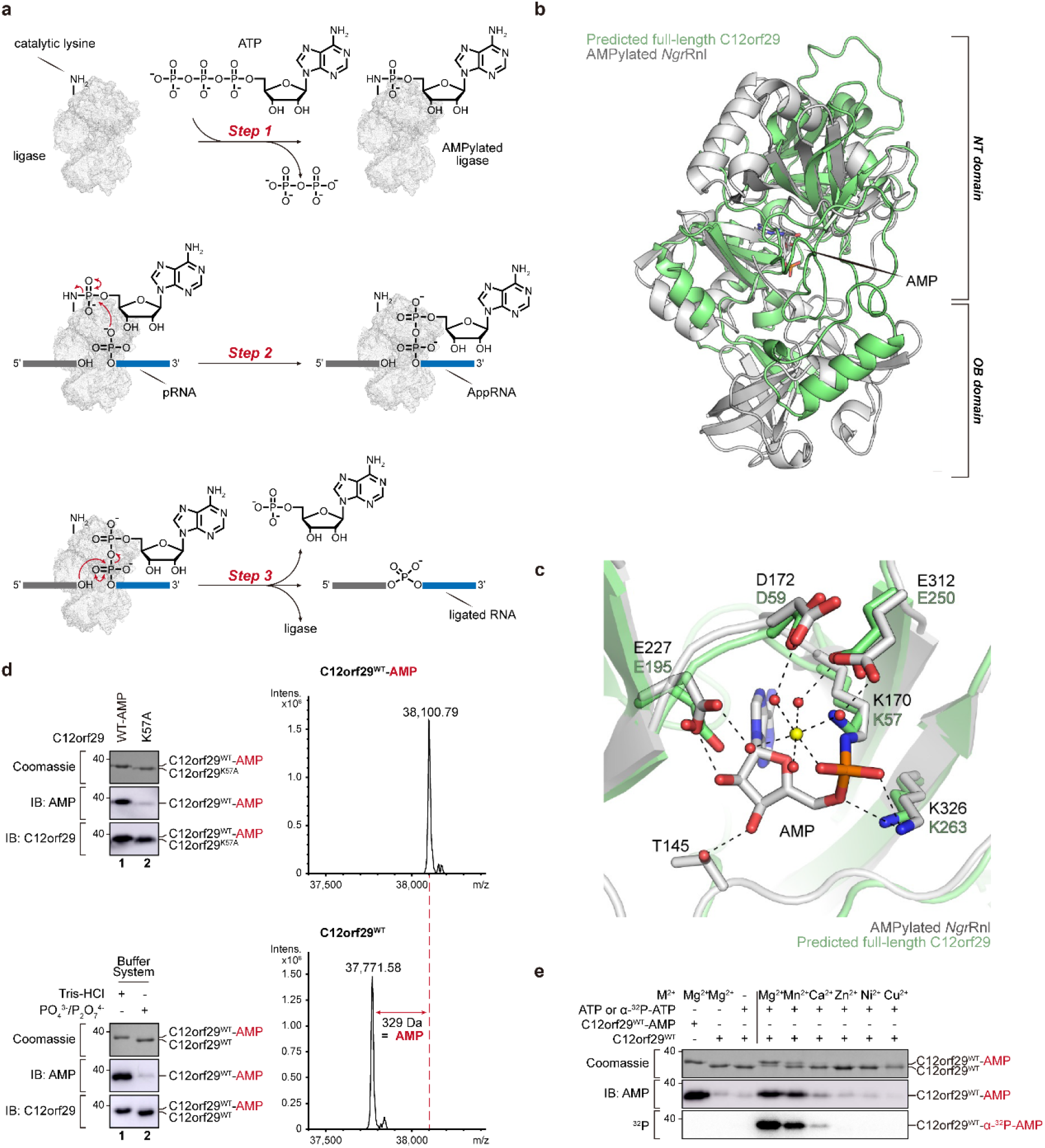
RNA ligase mechanism, structural prediction and auto-AMPylation activity of C12orf29. **a**, Schematic display of the three-step mechanism of RNA ligation by a 5’-3’ RNA ligase. In step 1, the ligase is auto-AMPylated on the catalytic lysine using ATP as the cosubstrate. In step 2, the AMP is transferred from the catalytic lysine to the 5’-PO_4_ end of RNA (pRNA), giving the RNA-adenylate intermediate (AppRNA). In step 3, the ligated RNA is obtained upon the attack of 3’-OH to the AppRNA in the presence of the ligase along with the liberation of AMP. **b**, Superimposition of the structure of C12orf29 predicted by AlphaFold (green)^14,15^ on the structure of *Ngr*Rnl (PDB ID: 5COT, grey). OB, oligonucleotide-binding. NT, nucleotidyltransferase. Structures were superposed in Coot^28^ using structural equivalent residues identified by the DALI webserver^16^. **c**, Enlarged view of the catalytic pocket of *Ngr*Rnl (PDB ID: 5COT, grey) and predicted C12orf29 (green). In the *Ngr*Rnl structure, AMP is covalently attached to the side chain of K170. D172, E227, and E312 bind Mn^2+^ via water-mediated contacts and K326 contacts the phosphate moiety of AMP. Corresponding residues in the putatively catalytic pocket of C12orf29 are indicated. Mn^2+^ and water molecules were depicted as yellow and red spheres, respectively. Atomic contacts were depicted as dashed lines. **d**, Top: Immunoblotting of C12orf29^WT^-AMP and C12orf29^K57A^ and MS analysis of C12orf29^WT^-AMP. C12orf29^K57A^ is deficient in auto-AMPylation activity. Mass spectra indicating a 329 Da increase in mass upon auto-AMPylation of C12orf29^WT^. C12orf29^WT^-AMP: calc. 38,100.34 Da, found 38,100.79 Da. C12orf29^WT^: calc. 37,771.29 Da, found 37,771.58 Da. Bottom: Preparation of C12orf29^WT^ and C12orf29^WT^-AMP with different buffer systems. **e**, Divalent metal ion dependency of auto-AMPylation activity of C12orf29^WT^.

The structural similarity to *Ngr*Rnl motivated us to investigate whether C12orf29 also possesses RNA ligase activity. Recently, AlphaFold^14^ structure predictions for almost all proteins of the human genome were made available at https://alphafold.ebi.ac.uk^15^. We submitted the AlphaFold structure prediction for C12orf29, which was expected to be of high quality (average pLDDT = 91.4), to the Dali server^16^, and compared it exhaustively to a representative subset of the Protein Data Bank. The results of structural superposition by Dali are highly significant and display RNA ligases as most similar structures (top hit, Z-score = 7.4 for *Ngr*Rnl), confirming our initial sequence-based suggestion for the functional assignment of C12orf29. In turn, the structure of C12orf29 predicted by AlphaFold was superimposed on *Ngr*Rnl (Fig. 2b, c and Supplementary Fig. 2). While an N-terminal oligonucleotide-binding (OB) domain as in *Ngr*Rnl was not observed for C12orf29, several secondary structure patterns in the predicted C12orf29 structure precisely overlapped with the C-terminal nucleotidyltransferase (NT) domain of *Ngr*Rnl (Fig. 2b, c). Of note, C12orf29 contains a lysine at position 57 within the conserved sequence motif KX(D/H/N)G that defines the superfamily of nucleotidyltransferases (Supplementary Fig. 2)^6^.

To elucidate potential enzymatic activities of C12orf29, we prepared the recombinant full-length wild-type (WT) protein. Surprisingly, LC-MS measurement of the protein purified from *Escherichia coli* (*E. coli*) revealed a molecular mass that is 329 Da higher than the calculated mass of 37,771 Da suggesting that C12orf29 is AMPylated (Fig. 2d). Indeed, AMPylation of C12orf29 was validated by Western blot (WB) analysis using a monoclonal antibody against AMPylation^17^ providing the first evidence that C12orf29 may act as an RNA ligase (Fig. 2d). Meanwhile, we also developed an expression and purification protocol that allowed the preparation of C12orf29 without AMPylation (Fig. 2d and see details in Supplementary Information). Interestingly, the AMPylated and the non-modified form of C12orf29 showed different migration behaviours in SDS-PAGE. Moreover, in line with our expectation, mutation of the critical lysine at position 57 to alanine (K57A) abolished AMPylation. With C12orf29^WT^ and C12orf29^WT^-AMP in hand, we further proved that C12orf29 was auto-AMPylated using ATP as a cosubstrate in the presence of Mg^2+^ as the most proficient cofactor (Fig. 2e).

Encouraged by the auto-AMPylation activity, we next investigated whether C12orf29 has 5’-3’ RNA ligase activity. We initially studied nick sealing activity as described for *Ngr*Rnl and used substrates that consists of an oligonucleotide splint bringing two oligonucleotides in a complex to form a nick to be ligated^15^ but did not detect any activity (Supplementary Fig. 3). In our further search for suitable substrates, we were guided by the following hypothesis: Compared to double-stranded RNA regions, the increased flexibility of single-stranded RNA (ssRNA) allows in-line attack of the 2’-OH group at the bridging phosphodiester bond^18^. This favours transesterification and in consequence RNA strand cleavage. We further reasoned that enzymes that repair and ultimately ligate such ssRNA strand breaks should be beneficial to a cell^4,5,19^. Therefore, we focused on ssRNA as substrates and studied oligonucleotide constructs that were composed of two strands and folded into an “open hairpin”. The 5’-terminus of one strand (17 nt) at the putative ligation site was labelled with ^32^P-phosphate while the other strand (10 nt) was unlabelled (Fig. 3a and Supplementary Fig. 4). Successful ligation would lead to 27 nt oligonucleotide with significantly altered migration detectable by denaturing PAGE analysis. Indeed, we found that C12orf29 was proficient in ligating the ssRNA overhangs to give a product that migrated at the expected length (Fig. 3a). Moreover, formation of cyclized RNA as well as AppRNA was also observed. Interestingly, the enzyme exhibited profound selectivity for RNA: RNA constructs were efficiently ligated within short time while the respective DNA constructs were not detectably converted under the very same conditions. We also investigated whether nucleotides other than ATP represented proficient cosubstrates and found that other adenosine nucleotides such as diadenosine tri- and tetraphosphate (Ap_3_A, Ap_4_A), and to a lesser extent even dATP are used as cosubstrates, while NAD^+^ is not used for promoting RNA ligation (Fig. 3b). Among the other nucleotides investigated, only GTP can promote the RNA ligation reaction (Fig. 3b). Additional kinetic investigations, however, revealed that the apparent catalytic efficiency (*k*_cat_/*K*_M_) of C12orf29 is about 29-fold higher with ATP than with GTP. This difference is mainly due to differences in *K*_M_ while the *k*_cat_ is similar for both nucleotides (Fig. 3c and Supplementary Fig. 5).

**Fig. 3.**
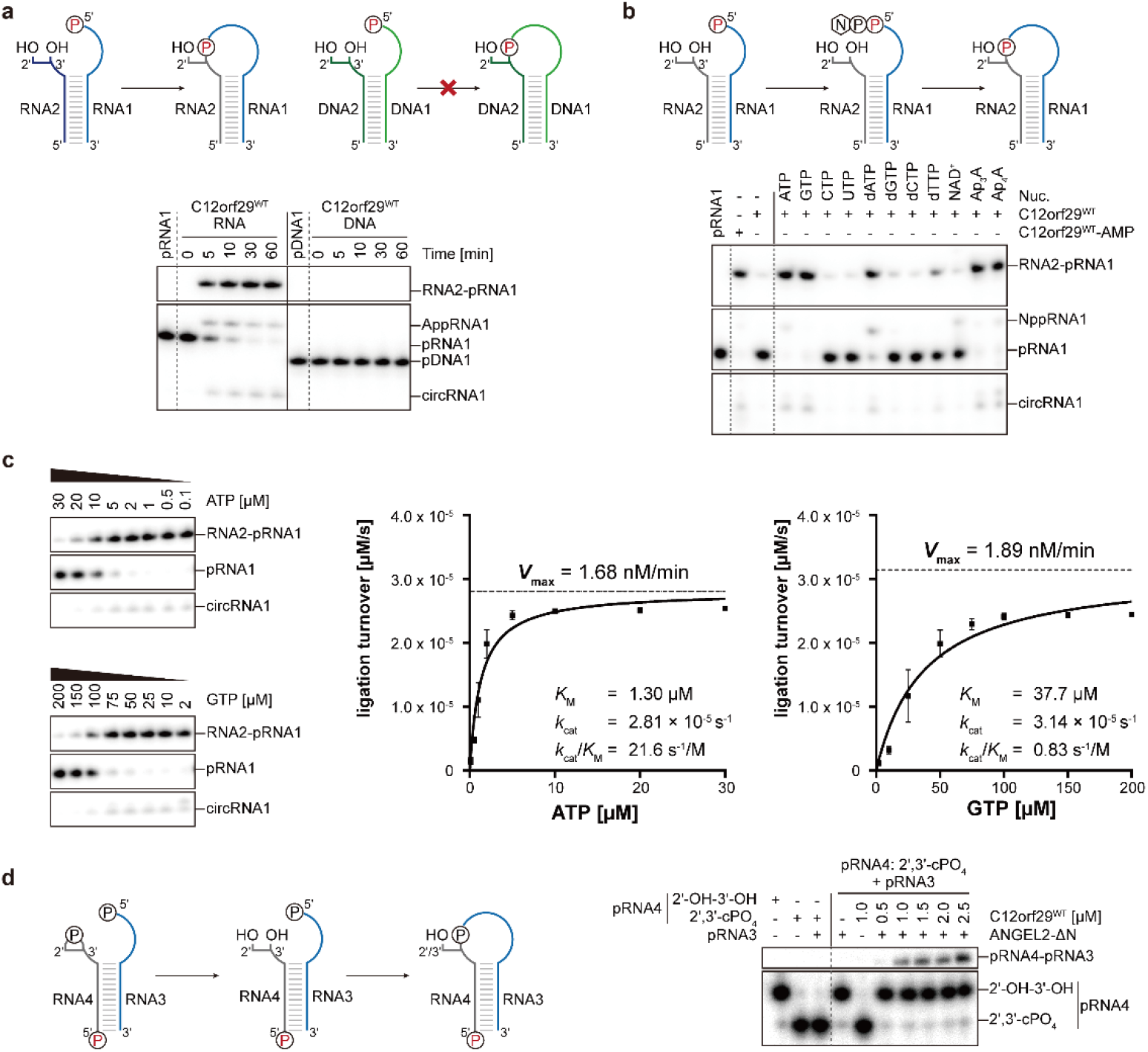
RNA ligase activity of C12orf29. RNA/DNA oligonucleotides are schematically depicted in blue/green, respectively. The ^32^P-labeled 5’-ends were depicted in red. The radioactive oligonucleotides were resolved by denaturing PAGE and analysed by phosphorimaging. **a**, (Top) Schematic display of two nucleic acid substrates used. (Bottom) PAGE analysis of reaction products after incubation with C12orf29 for various time points as indicated. Ligation was observed for 5’-phosphorylated RNA. **b**, Reaction of C12orf29 with indicated RNA constructs using various nucleotides. Besides ATP, C12orf29 is able to efficiently catalyse ligation by processing GTP, dATP, Ap_3_A and Ap_4_A. **c**, (Left) C12orf29 promoted ligation reaction at various concentrations of ATP and GTP. (Right) Michaelis-Menten fits to initial rates providing *K*_M_, *k*_cat_ and *k*_cat_/*K*_M_ for ATP and GTP. Plotted data represent the mean value ± SD for three biological replicates. **d**, (Left) Proposed ligation scheme starting from 2’,3’-cyclic phosphorylated RNA by the sequential action of ANGEL2-ΔN and C12orf29^WT^. (Right) PAGE analysis of the depicted reaction. All oligonucleotide sequences are provided in Supplementary Fig. 4.

Next, we investigated if 3’-termini bearing a 2’,3’-cyclic phosphate (cPO_4_) or a 2’-phosphate end can be used by C12orf29 in the ligation reaction. We found that both constructs were not processed by the protein (Supplementary Fig. 6). However, when the RNA substrate with a 2’,3’-cPO_4_ at the 3’-terminus was incubated in the presence of N-terminus-truncated ANGEL2 (ANGEL2-ΔN), an enzyme recently reported to cleave this phosphate from the 3’-terminus resulting in non-phosphorylated 3’-termini^20^, we observed C12orf29-dependent ligation to occur (Fig. 3d). Conclusively, these results show that non-phosphorylated 3’-RNA termini are essential for the ligase reaction.

In order to investigate the structural requirements for C12orf29-catalyzed ligation within the single-stranded regions, we designed and studied single RNA oligonucleotides modified with ^32^P-phosphate at the 5’ ends, which fold into structures bearing two single-stranded regions at the 5’ and 3’ ends (Fig. 4a, b and Supplementary Fig. 4). When varying the nucleotide composition at the ligation site we found that RNA bearing purines at the ligation site are most efficiently processed (Fig. 4a). Moreover, longer 5’-overhangs are also ligated more efficiently (Fig. 4b).

**Fig. 4.**
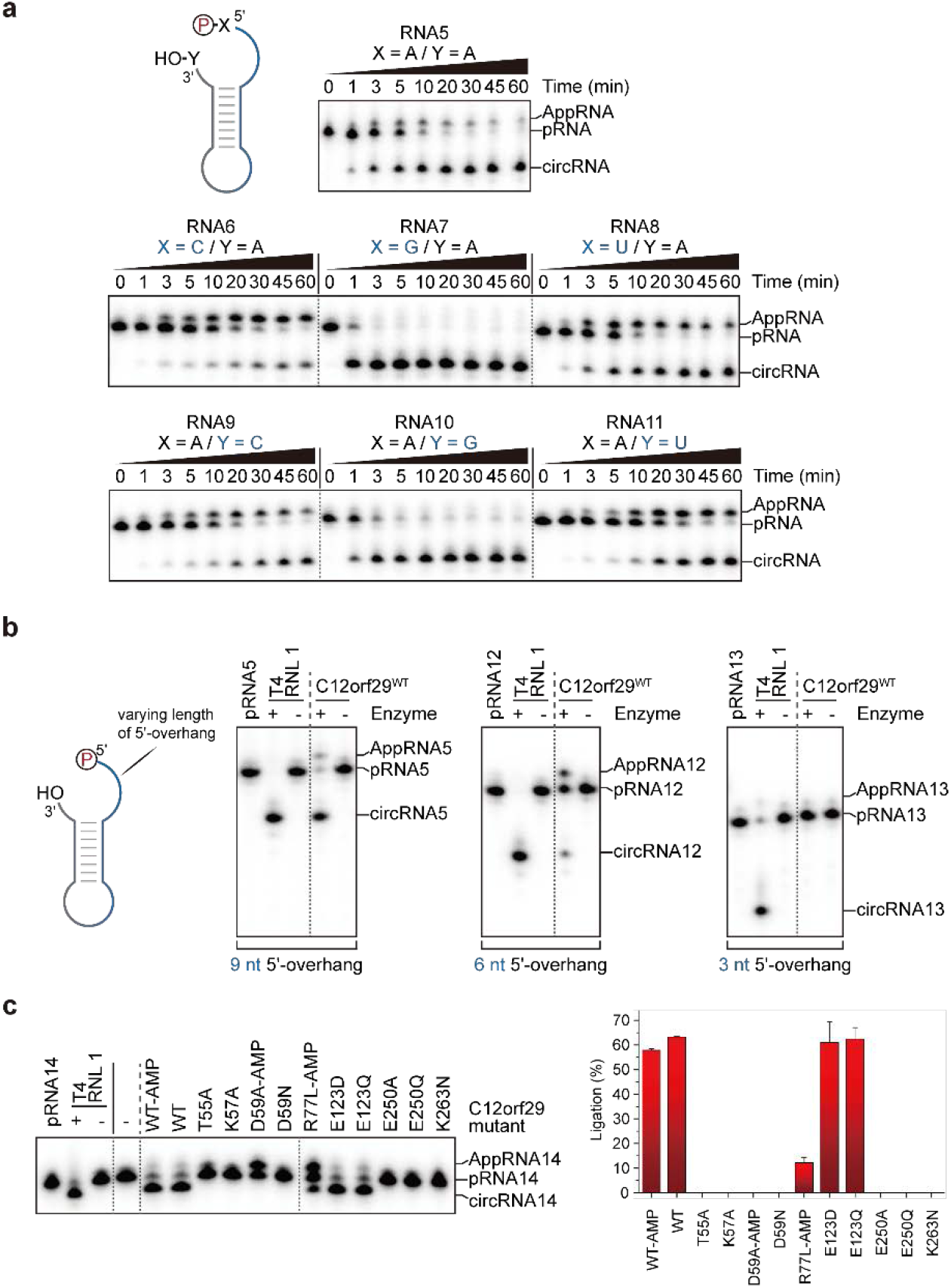
Substrate scope of C12orf29. RNA oligonucleotides that were used for intramolecular ligation resulting in cyclization are schematically depicted. The ^32^P-labeled 5’-ends were depicted in red. The radioactive oligonucleotides were resolved by denaturing PAGE and analysed by phosphorimaging. **a**, Investigation of the impact of the nucleobase composition at the 5’- and 3’-termini at the ligation site on the ligation efficiency of C12orf29. The depicted constructs were incubated under the same conditions for various time points. Most efficient ligation was observed when purines were at the ligation site. **b**, Investigation of the impact of the length of the 5’-overhang on the ligation efficiency of C12orf29. **c**, Impact of single site mutations on RNA ligase efficiency of C12orf29. Plotted data represent the mean value ± SD for three biological replicates. All oligonucleotide sequences are provided in Supplementary Fig. 4.

As shown in Fig. 2c and Supplementary Fig. 1 and 2, several conserved residues are proposed to be critical for the RNA ligase activity of C12orf29 since they make up the catalytic centre and are proposed to be involved e.g., in substrate and cofactor coordination. To examine the importance of these residues for RNA ligation as well as some residues that were reported to be mutated in cancers, a series of C12orf29 variants bearing mutations at the respective residues were prepared and tested. We found that several of these mutations abolish RNA ligation (Fig. 4c). Noteworthy, among the C12orf29 variants studied, the mutations D59N, R77L, E123D, and K263N were found in patients suffering from esophageal squamous cell carcinoma^21^, glioblastoma^22^, chronic lymphocytic leukemia^23^, and ovarian serous cystadenocarcinoma^24^, respectively. While E123D has no influence on the RNA ligase activity, D59N, R77L, and K263N impair RNA ligation.

### Knockout (KO) of C12orf29 increases cellular vulnerability towards reactive oxygen species

To gain insights into the function of C12orf29 in a cellular context, we generated human embryonic kidney (HEK293) cells, in which the *C12ORF29* gene (Supplementary Fig. 7) was knocked-out by CRISPR/Cas, and compared the properties of these cells with those of parental HEK293 cells expressing the WT enzyme. When cells were grown under physiological conditions, WT and KO cells have similar phenotypes with respect to cell growth, adherence, and appearance (Fig. 5a). However, when cells were treated with menadione, a molecule known to generate reactive oxygen species (ROS)-based cellular stress^25,26^, significant differences were observed (Fig. 5a). In comparison to WT cells, KO cells start to round up and to detach at lower concentrations of menadione. Moreover, KO cells are much more vulnerable to menadione so that cell viability is affected at lower concentration of menadione in comparison to WT cells (Fig. 5b). Next, we investigated whether this difference stems from a varying ROS formation in those cell lines. We found that KO cells die at much lower ROS levels than their WT counterpart (Fig. 5c). Therefore, we conclude that the observed increase in vulnerability towards ROS of the KO cells is not caused by difference of ROS formation in the cell lines.

**Fig. 5.**
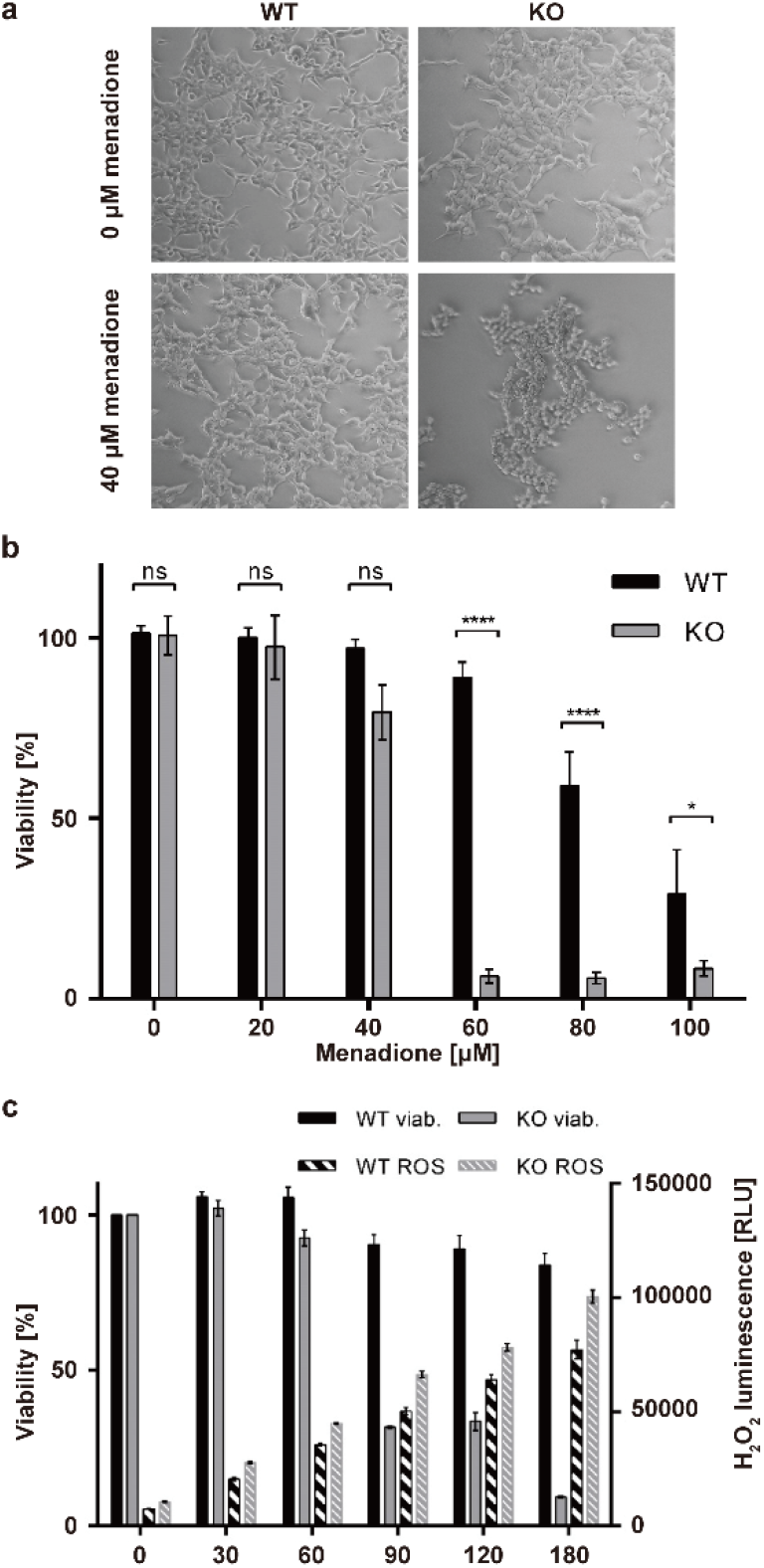
Effect of menadione-induced oxidative stress on WT and C12orf29-KO cells. **a**, Light microscopy of HEK293 WT and *C12ORF29*-KO cells. The cells were treated with 40 μM menadione for 3 h or only with the carrier as control (0 μM menadione). **b**, Cell viability of HEK293 WT and *C12ORF29*-KO cells after 3 h treatment with different menadione concentrations (n = 9, error bars represent the ± SD. Significance was calculated by two-way ANOVA with Sidak’s multiple comparisons test: ns = P > 0.05; * = P ≤ 0.05; **** = P ≤ 0.0001). **c**, Cell viability and corresponding H_2_O_2_ concentrations in HEK293 WT and *C12ORF29*-KO cells at different time points after treatment with 40 μM menadione (n = 3, error bars represent the ± SD). Viab = viability.

Next, we isolated and analysed the total RNA from the two cell lines treated with various concentrations of menadione (Fig. 6). Very prominent in this analysis are the signals of 28S and 18S rRNA. We found that the amount of RNA degraded depends on the concentration of menadione applied. Interestingly, while the signal for 18S rRNA remains almost constant even at higher menadione concentrations, 28S rRNA appears to be more vulnerable. More importantly, in KO cells 28S rRNA is already degraded at a much lower concentration of menadione than in WT cells. (Fig. 6 and Supplementary Fig. 8). Since the intracellular menadione-triggered ROS formation does not vary between both cell lines (Fig. 4c), we conclude that the observed increase in RNA decay in the KO cells is a result of the lack of C12orf29. Therefore, we conclude that C12orf29 is important in maintaining RNA integrity under stress conditions.

**Fig. 6.**
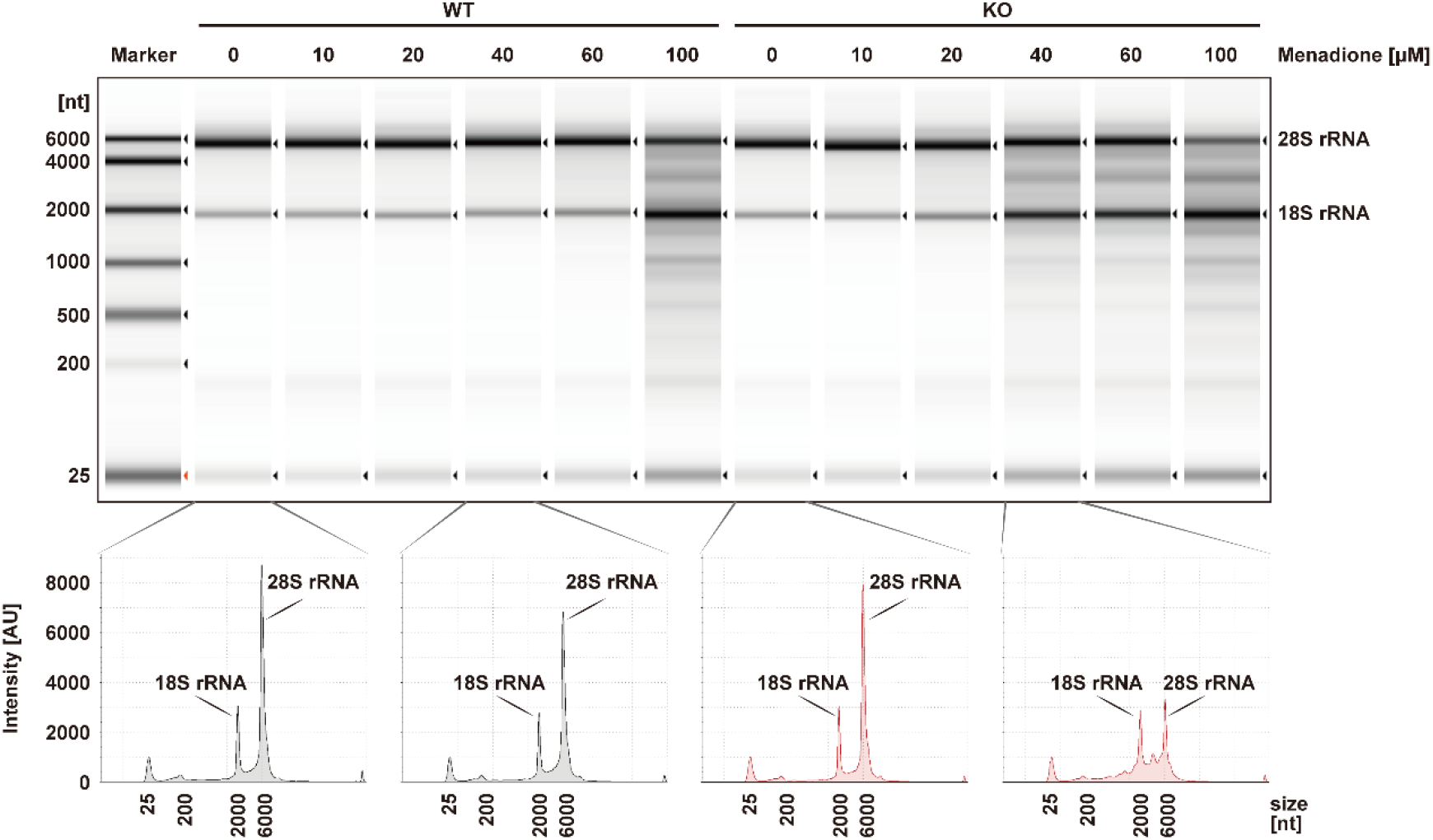
Analysis of total RNA from cell extracts of HEK293 WT and *C12ORF29*-KO cells treated with various concentrations of menadione. (Top) Cells (left: HEK293 WT cells; right: *C12ORF29*-KO HEK293 cells) were treated with different menadione concentrations for 3 h. In turn, total RNA was isolated and subsequently analysed by TapeStation. (Bottom) Electropherograms of the TapeStation analysis of total RNA from cell extracts of HEK293 WT (in black) and *C12ORF29*-KO (in red) cells.

## Discussion

This study reports on the discovery of human C12orf29 as the first 5’-3’ RNA ligase in higher eukaryotes. Our results elucidate the substrate spectrum of C12orf29 and show that the enzyme ligates RNA via successive protein and RNA AMPylation. Our studies show that HEK293 cells are more vulnerable to menadione-triggered ROS when *C12ORF29* is knocked out. We found decrease of resilience towards ROS in the KO cells with a simultaneous increase in RNA decay indicating that RNA ligation has an important role in stress-adaptive RNA repair.

Indeed, enzymes with 2’,3’-cyclic phosphatase (ANGEL2)^21^ and 5’-kinase (*Hs*CLP1)^27^ activities have been identified in humans and may work in concert with C12orf29 in the repair of RNA strands that result from transesterification. The data obtained indicates the importance of maintaining a functional RNA pool (ribonucleostasis) for cellular fitness. Since malfunction of RNA repair in humans is connected to the onset of several diseases such as neurodegeneration and cancer^5,19^ our findings may have implications for the development of new therapeutics. Our results demonstrate that the human protein C12orf29 is an RNA ligase and therefore we propose to name the protein Homo sapiens RNA ligase (*Hs*Rnl).

## Supporting information

Supplemental Figures, Methods, NMR data etc.

## Methods

### Chemical proteomics towards the identification of C12orf29

The synthesis of the probes is detailed in the Supplementary Information. In the chemical proteomics assay, 200 μM Ap_3_A, *C*2-eAp_3_A, or MilliQ^®^ H_2_O were incubated with 2.0 mg/mL H1299 or HEK293T cell lysates in 1x AMPylation buffer (20 mM HEPES pH 7.4, 100 mM NaCl, 5 mM MgCl_2_, and 1 mM DTT) at 37°C for 1 h in a total volume of 450 μL. The reaction was stopped by adding 1.8 mL pre-cold MeOH. The resulting mixture stood at -20 °C for 2 h to precipitate. Protein pellets were obtained after centrifugation at 14,000 x g for 10 min at 4 °C, which were dried for 5 min and reconstituted in 450 μL 1x resuspension buffer (50 mM triethanolamine pH 7.4, 150 mM NaCl, and 4% SDS). A master mix was prepared freshly with 0.5 mM N_3_-(Arg)PEG_3_-DB, 2.5 mM CuSO_4_, 0.25 mM TBTA, and 2.5 mM TCEP in 0.4x AMPylation buffer. 300 μL of the master mix was added to the pre-cold reaction mixture to yield 0.2 mM N_3_-(Arg)PEG_3_-DB, 1.0 mM CuSO_4_, 0.1 mM TBTA, and 1.0 mM TCEP in a total volume of 750 μL. The CuAAC was conducted at 25 °C for 1 h, which was quenched by adding 3 mL pre-cold acetone. The resulting mixture stood at -20 °C overnight to precipitate. Protein pellets were obtained after centrifugation at 14,000 x g for 10 min at 4 °C, which were washed with 300 μL cold MeOH trice and dried for 5 min. The pellets were reconstituted in 200 μL 1x PBS pH 7.4 supplemented with 4% SDS, followed by addition of 800 μL 1x PBS pH 7.4 and centrifugation at 13,400 rpm for 5 min at room temperature to remove any undissolved residue. The supernatant was incubated with high capacity streptavidin agarose beads in a bed volume of 25 μL at 25 °C for 15 min with an end-over-end rotator. The beads were pelleted by centrifugation at 1500 rpm for 2 min at room temperature, which were washed successively with 1x PBS pH 7.4 supplemented with 1% SDS (3 × 100 μL), washing buffer (8 × 100 μL, 1x PBS pH 7.4, 150 mM NaCl, 4 M urea, and 1% SDS), and 50 mM NH_4_HCO_3_ pH 7.8 (5 × 100 μL). The beads were treated with 0.8 mM biotin in 50 mM NH_4_HCO_3_ pH 7.8 supplemented with 0.1% RapiGest SF (3 × 50 μL) and incubated at 37°C for 10 min with shaking at 600 rpm to elute the AMPylated proteins. The elution fractions were kept on ice for downstream in-solution digestion (see below).

Prior to MS analysis, the elution fractions were treated with 5 mM DTT at 60 °C for 1 h. After cooling down, the mixture was treated with 50 mM 2-chloroacetamide at room temperature for 1 h in dark. In turn, the resulting mixture was incubated with 3 μg trypsin at 37 °C overnight. The digested protein mixture was added 3 μL TFA and incubated at 37 °C for 45 min to hydrolyse RapiGest SF. The resulting mixture was cleared by centrifugation at 13,400 rpm for 30 min at room temperature. The supernatant was transferred carefully and lyophilized overnight. Finally, the tryptic peptides were dissolved in 60 μL 0.1% TFA supplemented with 10 μL 10% TFA and desalted using U-C18 ZipTips. Tryptic peptides were separated on an EASY-nLC 1200 system (Thermo Scientific) at a flow rate of 300 nl/min using a 39 min gradient from 2.5% MeCN/0.1% formic acid to 32% MeCN/0.1% formic acid, 1 min to 75% MeCN/0.1% formic acid, followed by a 5 min washing step at 75% MeCN/0.1% formic acid. Mass spectra were recorded on a QExactive HF mass spectrometer (Thermo Scientific) operated in data dependent Top20 mode with dynamic exclusion set to 5 s. Full scan MS spectra were acquired at a resolution of 120,000 (at m/z 200) and scan range 350-1,600 m/z with an automatic gain control target value of 3e6 and a maximum injection time of 60 ms. Most intense precursors with charge states of 2-6 reaching a minimum automatic gain control target value of 1e3 were selected for MS/MS experiments. Normalized collision energy was set to 28. MS/MS spectra were collected at a resolution of 30,000 (at m/z 200), an automatic gain control target value of 1e5 and 100 ms maximum injection time. Each of the biological triplicates was measured as technical duplicates.

Raw files from LC-MS/MS measurements were analysed using MaxQuant (version 1.6.1.0) with match between runs and label-free quantification (LFQ) (minimum ratio count 1) enabled. The minimal peptide length was set to 5. For protein identification, the human reference proteome downloaded from the UniProt database (download date: 2018-02-22) and the integrated database of common contaminants were used. Data of different cell types were quantified in separate analysis. Further data processing was performed using Perseus software (version 1.6.1.3). Identified proteins were filtered for reverse hits, common contaminants and proteins that were only identified by site. LFQ intensities were log2 transformed, filtered to be detected in at least 4 out of 6 replicates and missing values were imputed from a normal distribution (width 0.3 and shift 1.8), based on the assumption that these proteins were below the detection limit. Significantly enriched proteins were identified by an ANOVA test (HEK293T cells: FDR = 0.001, S0 = 10; H1299 cells: FDR = 0.001, S0 = 6), normalized by Z-scoring and averaged.

Alternatively, the elution fractions were added 6x loading buffer and heated to 95 °C for denaturation and concentration. The resulting mixture was resolved by SDS-PAGE and subjected to immunoblotting.

### Expression and Purification of Recombinant Proteins

As for expression and purification of AMPylated C12orf29^WT^ and its variants, plasmid constructs pET15b-C12orf29^WT^ or plasmids containing inserts of C12orf29 variants were transformed in *E. coli* BL21 (DE3) competent cells, which were cultured in 50 mL LB medium containing 100 μg/mL carbenicillin at 37 °C, 180 rpm overnight. In turn, a defined volume of cell suspension was transferred to 1 L LB medium containing 100 μg/mL carbenicillin to reach OD_600_ = 0.1, followed by the incubation at 37 °C at 180 rpm until OD_600_ = 0.7. The mixture was cooled down on ice for 30 min and then incubated with 1.0 mM IPTG at 18 °C for 18 h at 180 rpm. Cells were harvested by centrifugation at 4,400 rpm for 30 min at 4 °C. The pellet was resuspended in 30 mL cold lysis buffer (50 mM Tris-HCl pH 8.0, 150 mM NaCl, 1.0 mM DTT, 0.1% (v/v) Triton X-100, 20 mM imidazole, 1 μg/mL aprotinin, 1 μg/mL leupeptin, and 1 mg/mL Pefabloc^®^ SC) and lysed by sonication on ice. The lysates were centrifuged at 40,000 x g for 30 min at 4 °C and filtered through 0.45 μm syringe filter. The N-terminal His_6_-tagged AMPylated C12orf29 was purified using a 5 mL HisTrap™ FF crude column (Buffer A: 50 mM Tris-HCl pH 8.0, 150 mM NaCl, 1.0 mM DTT, and 20 mM imidazole; Buffer B: 50 mM Tris-HCl pH 8.0, 150 mM NaCl, 1.0 mM DTT, and 500 mM imidazole). Fractions containing AMPylated His_6_-C12orf29 were pooled and dialyzed against a buffer containing 50 mM Tris-HCl pH 8.0, 100 mM NaCl, 5 U/mg thrombin, and 1 mM DTT at 4 °C overnight. The resulting solution was further purified by anion IEX on a 5 mL HiTrap™ Q HP column (Buffer A: 50 mM Tris-HCl pH 8.0, 100 mM NaCl, and 1.0 mM DTT; Buffer B: 50 mM Tris-HCl, pH 8.0, 1000 mM NaCl, and 1.0 mM DTT). Pure fractions were pooled, concentrated, and stored at -20 °C in a storage buffer containing 25 mM Tris-HCl pH 8.0, 100 mM NaCl, 1 mM DTT, and 50% (v/v) glycerol.

As for expression and purification of deAMPylated C12orf29^WT^ and its variants, plasmid constructs pET15b-C12orf29^WT^ or plasmids containing inserts of C12orf29 variants were transformed in *E. coli* BL21 (DE3) competent cells, which were cultured in 50 mL LB medium containing 100 μg/mL carbenicillin at 37 °C, 180 rpm overnight. In turn, a defined volume of cell suspension was transferred to 1 L LB medium containing 100 μg/mL carbenicillin to reach OD_600_ = 0.1, followed by the incubation at 37 °C at 180 rpm until OD_600_ = 0.7. The mixture was cooled down on ice for 30 min and then incubated with 1.0 mM IPTG at 18 °C for 18 h at 180 rpm. Cells were harvested by centrifugation at 4,400 rpm for 30 min at 4 °C. The pellet was resuspended in 30 mL cold lysis buffer (50 mM KH_2_PO_4_ pH 8.0, 10 mM Na_4_P_2_O_7_, 150 mM NaCl, 1.0 mM DTT, 0.1% (v/v) Triton X-100, 20 mM imidazole, 1 μg/mL aprotinin, 1 μg/mL leupeptin, and 1 mg/mL Pefabloc^®^ SC) and lysed by sonication on ice. The lysates were centrifuged at 40,000 x g for 30 min at 4 °C and filtered through 0.45 μm syringe filter. The N-terminal His_6_-tagged deAMPylated C12orf29 was purified using a 5 mL HisTrap™ FF crude column (Buffer A: 50 mM KH_2_PO_4_ pH 8.0, 10 mM Na_4_P_2_O_7_, 150 mM NaCl, 1.0 mM DTT, and 20 mM imidazole; Buffer B: 50 mM KH_2_PO_4_ pH 8.0, 10 mM Na_4_P_2_O_7_, 150 mM NaCl, 1.0 mM DTT, and 500 mM imidazole). Fractions containing His_6_-C12orf29 were pooled and dialyzed against a buffer containing 50 mM Tris-HCl pH 8.0, 100 mM NaCl, 5 U/mg thrombin, and 1 mM DTT at 4 °C overnight. The resulting solution was further purified by anion IEX on a 5 mL HiTrap™ Q HP column (Buffer A: 50 mM Tris-HCl pH 8.0, 100 mM NaCl, and 1.0 mM DTT; Buffer B: 50 mM Tris-HCl, pH 8.0, 1000 mM NaCl, and 1.0 mM DTT). Pure fractions were pooled, concentrated, and stored at -20 °C in a storage buffer containing 25 mM Tris-HCl pH 8.0, 100 mM NaCl, 1 mM DTT, and 50% (v/v) glycerol.

As for expression and purification of ANGEL2-ΔN, plasmid constructs pET15b-ANGEL2-ΔN were transformed in *E. coli* BL21 (DE3) cells, which were cultured in 50 mL LB medium containing 100 μg/mL carbenicillin at 37 °C, 180 rpm overnight. In turn, a defined volume of cell suspension was transferred to 1 L LB medium containing 100 μg/mL carbenicillin to reach OD_600_ = 0.1, followed by the incubation at 37 °C at 180 rpm until OD_600_ = 0.7. The expression was induced by the addition of 1.0 mM IPTG after cooling down the mixture on ice for 30 min. The mixture was then incubated at 18 °C for 18 h at 180 rpm before the cells were harvested by centrifugation at 4,400 rpm for 30 min at 4 °C. The pellet was resuspended in 30 mL cold lysis buffer (50 mM Tris-HCl pH 8.0, 100 mM KCl, 1 mg/mL lysozyme, 0.1% (v/v) Triton X-100, 1 mM DTT, 1 μg/mL aprotinin, 1 μg/mL leupeptin, and 1 mg/mL Pefabloc^®^ SC) on ice for 45 min and sonicated. The lysate was centrifuged at 40,000 x g for 30 min at 4 °C and filtered through 0.45 μm syringe filter. The His_6_-tagged ANGEL2-ΔN was purified using a 5 mL HisTrap™ FF crude column (Buffer A: 50 mM Tris-HCl pH 8.0, 100 mM KCl, 1.0 mM DTT, and 20 mM imidazole; Buffer B: 50 mM Tris-HCl pH 8.0, 100 mM KCl, 1.0 mM DTT, and 500 mM imidazole). Fractions containing His_6_-tagged ANGEL2-ΔN were pooled and dialyzed against a buffer containing 50 mM Tris-HCl pH 8.0, 100 mM KCl, 5 U/mg thrombin, and 1 mM DTT at 4 °C overnight. The resulting solution was further purified by anion IEX on a 5 mL HiTrap™ Q HP column (Buffer A: 50 mM Tris-HCl pH 8.0, 100 mM KCl, and 1.0 mM DTT; Buffer B: 50 mM Tris-HCl, pH 8.0, 1000 mM KCl, and 1.0 mM DTT). Pure fractions were pooled, concentrated, and stored at -20 °C in a storage buffer containing 25 mM Tris-HCl pH 8.0, 50 mM KCl, 1 mM DTT, and 50% (v/v) glycerol.

### Identification of the Intact Protein Mass

All samples were purified by UHPLC on a Dionex UltiMate3000 (Thermo Fisher Scientific, Germany) using an analytical Zorbax 300SB-C8 column (150 mm x 2.1 mm) with 3.5 μm silica as a stationary phase (Agilent, USA). Prior to purification, all samples were acidified with 10% TFA. Gradient elution (3 min at 0% Buffer B; in 19 min to 80% Buffer B; then in 8 min to 100% Buffer B) with Buffer A (0.02% TFA in water) and Buffer B (0.02% TFA in MeCN/water (80:20, v/v)) was performed at a flow rate of 300 μL/min. The signals were monitored by UV absorbance at 220 nm.

Intact proteins were then analysed by direct infusion on an amazon speed ETD mass spectrometer (Bruker Daltonics) with a flow rate of 4 μL/min. The mass spectrometric data were acquired for about 10 minutes and the final mass spectrum was averaged over the whole acquisition time. Mass spectrometric data were evaluated and deconvoluted using the Compass Data Analysis Version 4.4 (Bruker Daltonics) software.

### General Procedure of C12orf29 Auto-AMPylation Assays

Unless otherwise noted, the C12orf29 auto-AMPylation assays were performed as follow. 1.0 μM C12orf29^WT^-AMP, C12orf29^WT^ or its variants were incubated with 200 μM ATP or a mixture of ATP:α-^32^P-ATP (185 TBq/mmol, Hartmann Analytic, FP-307) in 9:1 ratio in 1x auto-AMPylation buffer (50 mM Tris-HCl pH 8.5, 5 mM MgCl_2_, and 1 mM DTT) at 37 °C for 30 min in a total volume of 18 μL. The reaction was stopped by transferring 15 μL reaction mixture to a pre-cold PCR tube containing 0.43 μL 0.5 M EDTA, 0.36 μL 2 mg/mL BSA, and 3.16 μL 6x loading buffer (50 mM Tris-HCl pH 6.8, 10% (v/v) glycerol, 2% (w/v) SDS, and 1% (v/v) β-mercaptoethanol). The resulting mixture was heated at 95 °C for 5 min. Samples were resolved by SDS-PAGE and analysed by Coomassie staining, autoradiographic imaging, or immunoblotting.

### Preparation of 5’ ^32^P-labeling of Oligonucleotides

Oligonucleotides (1.0 μM) were incubated with 15 units of T4 PNK (New England BioLabs, M0202S) and 200 μM 0.555 MBq γ-^32^P-ATP (185 TBq/mmol, Hartmann Analytic, SRP-401) in 1x T4 PNK reaction buffer at 37 °C for 1 h in a total volume of 15 μL. The reaction was stopped by heating to 95 °C for 2 min. The excess amount of γ-^32^P-ATP was removed by gel filtration using Sephadex™ G-10 resin to give 5’ ^32^P-labled oligonucleotides in a concentration of 1.0 μM. When labeling RNA oligos bearing 2’,3’-cPO_4_ or 2’-PO_4_-3’-OH on the 3’-ends, T4 PNK 3’ phosphatase minus (New England BioLabs, M0236S) was used.

### General Procedures of RNA Ligation Assays

As for general procedure of RNA ligation with C12orf29, unless otherwise noted, the RNA ligation with C12orf29 were performed as follow. 0.1 μM 5’ ^32^P-labeled oligonucleotide substrates were incubated with 1 μM C12orf29^WT^ or its variants and 200 μM ATP in 1x RNA ligation buffer (50 mM Tris-HOAc pH 7.0, 5 mM MgCl_2_, and 1 mM DTT) at 37 °C 1 h in a total volume of 10 μL. The reaction was quenched by adding 10 μL stopping solution (80% (v/v) formamide, 20 mM EDTA, 0.025% (w/v) bromophenol blue, and 0.025% (w/v) xylene cyanol) and heating at 95 °C for 2 min. 1 μL of the resulting mixture was further diluted to give 0.005 μM 5’ ^32^P-labeled oligonucleotides. Samples were resolved by urea-PAGE and analysed by autoradiographic imaging.

In the time course study, the reaction was performed in a total volume of 20 μL. Aliquots (1.5 μL) were taken at 0, 1, 3, 5, 10, 20, 30, 45, and 60 min following C12orf29^WT^ treatment and quenched by adding 1.5 μL 50 mM EDTA pH 8.0 solution. 1 μL of the resulting mixture was further diluted to give 0.005 μM 5’ ^32^P-labeled oligonucleotides, which was heated at 95 °C for 2 min. Samples were resolved by urea-PAGE and analysed by autoradiographic imaging.

### Kinetic Analysis of C12orf29-catalyzed RNA Ligation

Steady state kinetics of C12orf29 were measured at different ATP or GTP concentrations. Nucleotide concentrations used were 30, 20, 10, 5.0, 2.0, 1.0, 0.5 and 0.1 μM for ATP and 200, 150, 100, 75, 50, 25, 10 and 2.0 μM for GTP. 0.1 μM ^32^P-labeled RNA oligo1 and 0.5 μM RNA oligo2 were incubated with C12orf29^WT^ and the indicated concentrations of nucleotides in 1x RNA ligation buffer at 37 °C for 1 h in a total volume of 10 μL. The reaction was stopped by adding 150 μL stopping solution and heating at 95°C for 2 min. 1 μL of samples were resolved by urea-PAGE and analysed by autoradiographic imaging. Enzymatic turnover was quantified by Image Lab. The parameters *k*_cat_, *K*_M_, and *k*_cat_/*K*_M_ were determined from a non-linear regression fit/Michaelis-Menten curve of the initial rates at each substrate concentration using GraphPad Prism. Plotted data represent the median value ± SD for three biological replicates.

### RNA Ligation in the Presence of ANGEL2-ΔN with RNA Substrates Bearing 2’,3’-cPO_4_ on the 3’ Ends

5.0 μM of the 5’-OH RNA oligo3 substrate was incubated with 10 units of T4 PNK and 1.0 mM ATP in 1x T4 PNK reaction buffer at 37 °C for 1 h in a total volume of 50 μl. The reaction was stopped by heating to 95 °C for 2 min. The excess of ATP was removed by gel filtration to yield the respective 5’ phosphorylated RNA oligo3 in a concentration of 5.0 μM. 0.5 μM of the ^32^P-labeled RNA oligo4 was mixed with 0.6μM of the non-radioactively 5’-phosphorylated RNA oligo3 in MQ. The RNA strands were annealed using the annealing program described in general procedure.

0.1 μM of the annealed 5’ ^32^P-labeled RNA oligo4 and with 5’ non-radioactively phosphorylated RNA oligo3 complexes were incubated with different concentrations of C12orf29, 1 μM ANGEL2-ΔN, and 2.0 mM ATP in 1x RNA ligation buffer at 37 °C for 1 h in a total volume of 10 μL. The reaction was quenched by adding 95 μL stopping solution to 5 μL of the sample and heating to 95 °C for 2 min. The samples were then resolved by urea-PAGE (12%) and analysed by autoradiographic imaging.

### Light-microscopy of Menadione-treated HEK293 Cells

1.2 × 10^6^ cells were seeded in 4 mL DMEM GlutaMAX™ medium (Gibco™, Thermo Fisher) supplemented with 10% (v/v) FCS on 6 cm cell culture dishes (Sarstedt). After 48 h, the cells were treated directly with either 4 μL EtOH (control, no menadione) or with 4 μL of 40 mM menadione in EtOH (final concentration = 40 μM menadione). After 3 h, pictures were taken with a light microscope using a 5x objective.

### Cell Viability Assay of HEK293 Cells Treated with Different Concentrations of Menadione

The CellTiter-Glo^®^ Luminescent Cell Viability Assay (Promega) was used according to the manufacturer’s instruction. 4.0 × 10^4^ cells per well were seeded one day before the menadione treatment in 90 μL DMEM GlutaMAX™ medium with 10% FCS in a 96-well plate (Sarstedt). The plate was incubated at 37 °C, 5% CO_2_, and 100% humidity for 24 h.

1,000x menadione stock solutions were prepared in ethanol and frozen in aliquots. On the day of the treatment, aliquots were thawed and diluted 1/100 in MilliQ^®^ H_2_O. The cells were treated with 10 μL of different concentrations of menadione in MilliQ^®^ H_2_O. The plate was incubated at 37 °C, 5% CO_2_, 100% humidity. After 3 h, the cells were equilibrated to RT (15 min) and 100 μL of CellTiter-Glo^®^ reagent was added per well and mixed thoroughly. The plate was incubated for 10 min on a shaker. 100 μL of the solution was transferred to a black 96-well plate and luminescence read out was performed with a plate reader (PerkinElmer Victor3™ Multilabel Counter 1420).

The luminescence values of the treated cells were compared to the value of the cells of the control treatment, which were treated with 0 μM menadione (equals 100% viability), to give the calculated cell viability.

### RNA Integrity Analysis of HEK293 WT and KO Cells Treated with Different Concentrations of Menadione

5.0 × 10^5^ cells were seeded in 2 mL DMEM GlutaMAX™ medium with 10% FCS in a 6-well plate (Sarstedt) 48 h prior to the experiment. 1,000x menadione stock solutions were prepared in ethanol and frozen in aliquots. 2 μL of the respective menadione stock was given to the cells resulting in the final desired menadione concentration for the treatment. After 3 h the cells were scraped down in the present medium and centrifuged (500 × g, 5 min, 4 °C). The cell pellet was washed with 1 mL ice-cold PBS and centrifuged again (500 × g, 5 min, 4 °C). The RNA was then extracted from the pellet using Quick-RNA Miniprep Kit (Zymo Research) with the provided in-column DNase I digest. The resulting RNA was analysed using Agilent 4150 TapeStation System.

### Cell Viability Assay in Combination with ROS assay

The ROS-Glo™ H_2_O_2_ Assay (Promega) was used according to the manufacturer’s instruction. 4.0 × 10^4^ cells per well were seeded one day before the menadione treatment in 70 μL DMEM GlutaMAX™ medium with 10% FCS in a 96-well plate (Sarstedt). The plate was incubated at 37 °C, 5% CO_2_, 100% humidity for 24 h. 3 h before the final readout, 20 μL of the provided H_2_O_2_ substrate in H_2_O_2_ substrate dilution buffer was directly added to the cells in their growing medium (final concentration was 25 μM). Afterwards, 10 μL of menadione stock solution in EtOH was added to give the desired end concentration in a final volume of 100 μL. The cells were incubated for the desired stress time at 37 °C. The cells were equilibrated at RT for 15 min.

A. For detection of ROS, 50 μL of the present medium of each well was transferred in a new, black 96-well plate and 50 μL of the provided ROS detection solution was added. After 20 min incubation at RT, the luminescence was measured using a plate reader (PerkinElmer Victor3™ Multilabel Counter 1420).
B. For the cell viability assay, 50 μL of CellTiter-Glo^®^ reagent was added to the remaining 50 μL medium in the 96-well plate and mixed thoroughly. The plate was incubated for 10 min on a shaker. 100 μL of the solution was transferred to a black 96-well plate and the luminescence was measured using a plate reader (PerkinElmer Victor3™ Multilabel Counter 1420).

## Acknowledgments

We thank A. Marquardt and A. Sladewska-Marquardt (Proteomics Center, University of Konstanz) for assistance in mass spectrometry. Y.Y., F.M.S., L.A.S., L.B.H., M.F., and E.H. thank the Konstanz Research School Chemical Biology for support. We thank D. Höpfner and A. Itzen, UKE Hamburg, for the provision of the anti-AMPylation antibody. A.M. acknowledges the European Research Council for funding of this project (ERC AdG AMP-Alarm). F.S. acknowledges the Deutsche Forschungsgemeinschaft for funding (STE 2517/5-1). O.P.S. acknowledges support by the Alexander von Humboldt-Foundation.

## Author contributions

Y.Y., F.M.S., L.A.S., O.P.S. and A.M. conceived the study and experimental approach; Y.Y., F.M.S., L.A.S., O.P.S. generated C12orf29 and its mutants, performed AMPylation and RNA ligation experiments, Y.Y. conducted the modelling and MS analysis. F.M.S. conducted cellular experiments. L.B.H. and M.F. provide expert expertise in PAGE analysis and AMPylation reactions, respectively. M.S., E.H., F.S., K.D. provided expert expertise on cellular studies, mass spectrometry and modelling, respectively. Y.Y., F.M.S., L.A.S., O.P.S. and A.M. analysed the data, A.M. wrote the manuscript with input from all authors.

