## Supplemental Figures, Methods, NMR data etc. for "A human RNA ligase that operates via auto- and RNA-AMPylation"

### Supplementary Information

**Table of Contents**

Chemical Proteomics……………………………………………………………...S3
Plasmid Construction……………………………………………………………...S5
Site-directed Mutagenesis…………………………………………………………S5
Expression and Purification of Recombinant Proteins………………………………..S5
Identification of the intact Protein Mass……………………………………………...S9
Immunoblotting……………………………………………………………………S9
General Procedure of C12orf29 Auto-AMPylation Assays………………….………S9
Preparation of 5' $^{32}$P-labeling of Oligonucleotides…………………………………S10
Preparation of Ligase Substrates…………………………………………...……S10
General Procedures of RNA Ligation Assays……………………………………...S11
Nucleotide Screening in RNA Ligation Assay (Fig. 2b)…………………………….S13
Autoradiographic Analysis…………………………………………………….…S13
Kinetic Analysis of C12orf29-catalyzed RNA Ligation (Fig. 2c)………………...…S14
RNA Ligation in the Presence of ANGEL2-ΔN with RNA
Substrates Bearing 2',3'-cPO4…………………………………………………….S14
Mutational Effects on RNA Ligation……………………………………………...…S14
Preparation of *C12ORF29* Knockout HEK293 Cell Lines…………………….…S15
Light-microscopy of Menadione-treated HEK293 Cells……………………………S15
Cell Viability Assay of HEK293 Cells Treated with Different
Concentrations of Menadione…………………………………………………….S15
RNA Integrity Analysis of HEK293 WT and KO Cells
Treated with Different Concentrations of Menadione….……………………………S16
Cell Viability Assay in Combination with ROS assay……………………………….S16
Supplementary Fig. 1-11…………………………………………………….S17-S27
Synthesis and analysis of chemical probes…………………………………S28-S36
References……………………………………………………………………..S37
NMR spectra. ……………………………………………………….S38-S47

The gene fragments of C12orf29$^{WT}$, AtRNL-CPD (667-1104), ANGEL2-ΔN (119-544), bearing 5'-NdeI and 3'-XhoI restriction cleavage sites were synthesized by Integrated DNA Technologies. The gene fragments were ligated into pJET1.2/blunt vector using CloneJET PCR Cloning Kit for amplification. The NdeI and XhoI restriction digested gene fragments and pET15b vector were isolated and ligated.

**Site-directed Mutagenesis**

50 ng plasmid DNA templates were mixed with 0.4 μM forward primers and 0.4 μM reverse primers in 1x Pfu reaction buffer with 0.4 mM dNTPs each and 2.5 units of Pfu Turbo DNA polymerase in a total volume of 50 μL. The reaction mixture was applied for PCR by the following program:

| Step | Conditions | |
|---|---|---|
| Initial denaturation | 95 °C, 2 min | |
| Denaturation | 95 °C, 15 s | 20 cycles |
| Primer annealing | 55 °C, 30 s | |
| Extension | 68 °C, 10 min | |
| Elongation | 68 °C, 12 min | |
| Stopping | 4 °C, pause | |

Afterwards, 10 μL of PCR reaction mixture was incubated with 40 units of DpnI in 1x CutSmart® buffer for 30 min at 37 °C in a total volume of 50 μL. 10 μL of the resulting mixture was transformed into NEB Turbo chemically competent *E. coli* cells and plasmids were isolated for analysis.

*Expression and Purification of Plant tRNA Ligase AtRNL*

Plasmid constructs pET28a-At-TRL (Addgene: #32242) were transformed in *E. coli* BL21 (DE3) RIL competent cells, which were cultured in 50 mL LB medium containing 50 μg/mL kanamycin at 37 $^{o}$C, 180 rpm overnight. In turn, a defined volume of cell suspension was transferred to 1 L LB medium containing 50 μg/mL kanamycin to reach $OD_{600}$ = 0.1, followed by the incubation at 37 $^{o}$C at 180 rpm until $OD_{600}$ = 0.7. The mixture was cooled down on ice for 30 min and then incubated with 0.4 mM IPTG and 2% (v/v) ethanol at 17 $^{o}$C for 20 h at 180 rpm. Cells were harvested by centrifugation at 4,400 rpm for 30 min at 4 $^{o}$C. The pellet was resuspended in 30 mL cold lysis buffer (50 mM Tris-HCl pH 7.5, 250 mM NaCl, 10% (v/v) glycerol, 0.1% (v/v) Triton-X100, 20 mM imidazole, 1 μg/mL aprotinin, 1 μg/mL leupeptin, and 1 mg/mL Pefabloc® SC) and lysed by sonication. The lysates were centrifuged at 40,000 x g for 30 min at 4 $^{o}$C and filtered through 0.45 μm syringe filter. The $His_6$-tagged AtRNL was purified using a 5 mL HisTrap™ FF crude column (Buffer A: 50 mM Tris-HCl pH 7.5, 250 mM NaCl, 10% (v/v) glycerol, and 20 mM imidazole; Buffer B: 50 mM Tris-HCl pH 7.5, 250 mM NaCl, 10% (v/v) glycerol, and 500 mM imidazole). Fractions containing $His_6$-tagged AtRNL were pooled and dialyzed against a buffer containing 50 mM Tris-HCl pH 7.5, 250 mM NaCl, 5 U/mg thrombin, 10% glycerol, and 20 mM imidazole at 4 $^{o}$C overnight. The resulting solution was purified again using a 5 mL HisTrap™ FF crude column (Buffer A: 50 mM Tris-HCl pH 7.5, 250 mM NaCl, 10% (v/v) glycerol, and 20 mM imidazole; Buffer B: 50 mM Tris-HCl pH 7.5, 250 mM NaCl, 10% (v/v) glycerol, and 500 mM imidazole). The AtRNL were recovered in the flow-through, concentrated, and stored at -20 $^{o}$C in a storage buffer containing 25 mM Tris-HCl pH 8.0, 100 mM NaCl, and 50% (v/v) glycerol.

*Expression and Purification of the Cyclic Phosphodiesterase Domain of Plant tRNA Ligase AtRNL (AtRNL-CPD)*

Plasmid constructs pET15b-AtRNL-CPD were transformed in *E. coli* BL21 (DE3) competent cells, which were cultured in 50 mL LB medium containing 100 μg/mL carbenicillin at 37 $^{o}$C, 180 rpm overnight. In turn, a defined volume of cell suspension was transferred to 1 L LB medium containing 100 μg/mL carbenicillin to reach $OD_{600}$ = 0.1, followed by the incubation at 37 $^{o}$C at 180 rpm until $OD_{600}$ = 1.0. The mixture was cooled down on ice for 30 min and then incubated with 1.0 mM IPTG and 2% (v/v) ethanol at 20 $^{o}$C for 18 h at 180 rpm. Cells were harvested by centrifugation at 4,400 rpm for 30 min at 4 $^{o}$C. The pellet was resuspended in 30 mL cold lysis buffer (50 mM Tris-HCl pH 8.0, 200 mM NaCl, 10% (v/v) glycerol, 0.2% (v/v) Triton X-100, 20 mM imidazole, 1 μg/mL aprotinin, 1 μg/mL leupeptin, and 1 mg/mL Pefabloc® SC) and lysed by sonication. The lysates were centrifuged at 40,000 x g for 30 min at 4 °C and filtered through 0.45 μm syringe filter. The $His_6$-tagged AtRNL-CPD was purified using a 5 mL HisTrap™ FF crude column (Buffer A: 50 mM Tris-HCl pH 8.0, 200 mM NaCl, 10% (v/v) glycerol, and 20 mM imidazole; Buffer B: 50 mM Tris-HCl pH 8.0, 200 mM NaCl, 10% (v/v) glycerol, and 500 mM imidazole). Fractions containing $His_6$-tagged AtRNL were pooled and dialyzed against a buffer containing 50 mM Tris-HCl pH 8.0, 200 mM NaCl, 5 U/mg thrombin, 10% glycerol, and 20 mM imidazole at 4 °C overnight. The resulting solution was purified again using a 5 mL HisTrap™ FF crude column (Buffer A: 50 mM Tris-HCl pH 8.0, 200 mM NaCl, 10% (v/v) glycerol, and 20 mM imidazole; Buffer B: 50 mM Tris-HCl pH 8.0, 200 mM NaCl, 10% (v/v) glycerol, and 500 mM imidazole). The AtRNL-CPD were recovered in the flow-through, concentrated, and stored at -20 °C in a storage buffer containing 25 mM Tris-HCl pH 8.0, 100 mM NaCl, and 50% (v/v) glycerol.

**Immunoblotting**

Proteins were resolved by SDS-PAGE, transferred onto the MeOH-activated 0.2 uM PVDF membrane (Amersham$^{TM}$), and probed with anti-C12orf29 (Santa Cruz Biotechnology, sc-390730), anti-p150 (Bioscience, 610474) and anti-AMPylation antibodies[17] kindly provided by Itzen Aymelt and colleagues, UKE, University of Hamburg. Blots were developed using Pierce$^{TM}$ ECL Western Blotting Substrate (Thermo Fisher) or Pierece$^{TM}$ Fast Western Kit, SuperSignal$^{TM}$ West Pico (Thermo Fisher).

**General Procedure of C12orf29 Auto-AMPylation Assays**

Unless otherwise noted, the C12orf29 auto-AMPylation assays were performed as follow. 1.0 µM C12orf29$^{WT}$-AMP, C12orf29$^{WT}$ or its variants were incubated with 200 µM ATP or a mixture of ATP:α-$^{32}$P-ATP (185 TBq/mmol, Hartmann Analytic, FP-307) in 9:1 ratio in 1x auto-AMPylation buffer (50 mM Tris-HCl pH 8.5, 5 mM $MgCl_2$, and 1 mM DTT) at 37 $^{o}$C for 30 min in a total volume of 18 µL. The reaction was stopped by transferring 15 µL reaction mixture to a pre-cold PCR tube containing 0.43 µL 0.5 M EDTA, 0.36 µL 2 mg/mL BSA, and 3.16 µL 6x loading buffer (50 mM Tris-HCl pH 6.8, 10% (v/v) glycerol, 2% (w/v) SDS, and 1% (v/v) β-mercaptoethanol). The resulting mixture was heated at 95 $^{o}$C for 5 min. Samples were resolved by SDS-PAGE and analysed by Coomassie staining, autoradiographic imaging, or immunoblotting.

**Preparation of 5' $^{32}$P-labeling of Oligonucleotides**

Oligonucleotides (1.0 µM) were incubated with 15 units of T4 PNK (New England BioLabs, M0202S) and 200 µM 0.555 MBq γ-$^{32}$P-ATP (185 TBq/mmol, Hartmann Analytic, SRP-401) in 1x T4 PNK reaction buffer at 37 $^{o}$C for 1 h in a total volume of 15 µL. The reaction was stopped by heating to 95 $^{o}$C for 2 min. The excess amount of γ-$^{32}$P-ATP was removed by gel filtration using Sephadex™ G-10 resin to give 5' $^{32}$P-labled oligonucleotides in a concentration of 1.0 µM. When labeling RNA oligos bearing 2',3'-c$PO_4$ or 2'-$PO_4$-3'-OH on the 3'-ends, T4 PNK 3' phosphatase minus (New England BioLabs, M0236S) was used.

**Preparation of Ligase Substrates**

*Preparation of Nicked RNA/DNA Duplexes*

0.5 µM 5' $^{32}$P-labeled RNA/DNA oligonucleotides were mixed with 1.0 µM DNA splint oligo and 2.5 µM of the second RNA/DNA oligo in 1x annealing buffer (10 mM Tris-HCl pH 6.8 and 200 mM NaCl) (Sequences are shown in fig. S4). The mixture was prepared freshly and applied for annealing by the following program:

| Temperature | Time |
|---|---|
| 65 $^{o}$C | 10 min |
| 37 $^{o}$C | 15 min |
| 22 $^{o}$C | 30 min |
| 4 $^{o}$C | pause |

*Preparation of RNA Oligo4 Bearing 2'-$PO_4$-3'-OH on the 3' Ends*

0.5 µM RNA oligo4 bearing 2',3'-c$PO_4$ were incubated with 2.0 µM AtRNL-CPD in 50 mM Tris-HCl pH 7.5, 10 mM $MgCl_2$, and 2 mM DTT in a total volume of 4.0 mL. The reaction was split in a 96-well plate and incubated at 37 $^{o}$C for 1.5 h. Each reaction was stopped by adding 5 µL pre-cold 50 mM EDTA pH 8.0 solution and heating at 95 $^{o}$C for 2 min. All resulting mixtures were combined and purified with Oligo Clean & Concentrator Kit (Zymo Research).

*Preparation of Broken RNA Hairpins*

For preparing the complexes with 5'-$PO_4$ and 2'-OH-3'-OH overhangs, 0.5 µM 5' $^{32}$P-labeled RNA were mixed with 2.5 µM RNA that donate 2'-OH-3'-OH ends in MilliQ® $H_2O$. The mixture was prepared freshly and applied for the annealing program shown below.

For preparing the complexes with 5'-$PO_4$ and 2'-$PO_4$-3'-OH overhangs, 0.5 μM 5' $^{32}$P-labeled RNA were mixed with 2.5 μM RNA that donate 2'-$PO_4$-3'-OH ends in MilliQ® $H_2O$. The mixture was prepared freshly and applied for the annealing program shown below.

For preparing the complexes with 5'-OH and 2',3'-c$PO_4$ overhangs, 0.5 μM 5' $^{32}$P-labeled RNA oligo4 bearing 2',3'-c$PO_4$ ends were mixed with 2.5 μM RNA that donate 5'-OH ends in MilliQ® $H_2O$. The mixture was prepared freshly and applied for the annealing program shown below.

For preparing the complexes with 5'-$PO_4$ and 2',3'-c$PO_4$ overhangs, 0.5 μM 5' $^{32}$P-labeled RNA oligo4 were mixed with 0.6 μM RNA oligo3 that donate 5'-$PO_4$ ends in MilliQ® $H_2O$. The mixture was prepared freshly and applied for the annealing program shown below.

The annealing step followed program:

| Temperature | Time |
|---|---|
| 95 °C | 2 min |
| 65 °C | 30 s |
| 55 °C | 30 s |
| 45 °C | 30 s |
| 25 °C | 30 s |
| 15 °C | 30 s |
| 10 °C | 30 s |
| 4 °C | pause |

**General Procedures of RNA Ligation Assays**

*General Procedure of RNA Ligation with C12orf29*

Unless otherwise noted, the RNA ligation with C12orf29 were performed as follow. 0.1 μM 5' $^{32}$P-labeled oligonucleotide substrates were incubated with 1 μM C12orf29$^{WT}$ or its variants and 200 μM ATP in 1x RNA ligation buffer (50 mM Tris-HOAc pH 7.0, 5 mM $MgCl_2$, and 1 mM DTT) at 37 °C 1 h in a total volume of 10 μL. The reaction was quenched by adding 10 μL stopping solution (80% (v/v) formamide, 20 mM EDTA, 0.025% (w/v) bromophenol blue, and 0.025% (w/v) xylene cyanol) and heating at 95 °C for 2 min. 1 μL of the resulting mixture was further diluted to give 0.005 μM 5' $^{32}$P-labeled oligonucleotides. Samples were resolved by urea-PAGE and analysed by autoradiographic imaging.

In the time course study, the reaction was performed in a total volume of 20 μL. Aliquots (1.5 μL) were taken at 0, 1, 3, 5, 10, 20, 30, 45, and 60 min following C12orf29$^{WT}$ treatment and quenched by adding 1.5 μL 50 mM EDTA pH 8.0 solution. 1 μL of the resulting mixture was further diluted to give 0.005 μM 5' $^{32}$P-labeled oligonucleotides, which was heated at 95 °C for 2 min. Samples were resolved by urea-PAGE and analysed by autoradiographic imaging.

*General Procedure of RNA Ligation with T4 RNA Ligase 1*

0.1 μM 5' $^{32}$P-labeled oligonucleotide substrates were incubated with 5 units of T4 RNA ligase 1 and 1.0 mM ATP in 1x T4 RNA ligase reaction buffer with 20% PEG8000 at 25 °C for 2 h in a total volume of 10 μL. The reaction was quenched by adding 10 μL stopping solution and heating at 95 °C for 2 min. 1 μL of the resulting mixture was further diluted to give 0.005 μM 5' $^{32}$P-labeled oligonucleotides. Samples were resolved by urea-PAGE and analysed by autoradiographic imaging.

*General Procedure of RNA Ligation with T4 RNA Ligase 2*

0.1 μM 5' $^{32}$P-labeled oligonucleotide substrates were incubated with 5 units of T4 RNA ligase 2 in 1x T4 Rnl2 reaction buffer with 10 mM $MgCl_2$ at 37 °C 1 h in a total volume of 10 μL. The reaction was quenched by adding 10 μL stopping solution and heating at 95 °C for 2 min. 1 μL of the resulting mixture was further diluted to give 0.005 μM 5' $^{32}$P-labeled oligonucleotides. Samples were resolved by urea-PAGE and analysed by autoradiographic imaging.

*General Procedure of RNA Ligation with AtRNL*

0.1 μM 5' $^{32}$P-labeled oligonucleotide substrates were incubated with 5 μM AtRNL and 100 μM ATP in 1x RNA ligation buffer (50 mM Tris-HOAc pH 7.0, 5 mM $MgCl_2$, and 1 mM DTT) at 37 °C 1 h in a total volume of 10 μL. The reaction was quenched by adding 10 μL stopping solution and heating at 95 °C for 2 min. 1 μL of the resulting mixture was further diluted to give 0.005 μM 5' $^{32}$P-labeled oligonucleotides. Samples were resolved by urea-PAGE and analysed by autoradiographic imaging.

*General Procedure of RNA Ligation with RtcB*

0.1 μM 5' $^{32}$P-labeled oligonucleotide substrates were incubated with 1 μM RtcB and 100 μM GTP in 1x RtcB reaction buffer with 1 mM $MnCl_2$ at 37 °C 1 h in a total volume of 10 μL. The reaction was quenched by adding 10 μL stopping solution and heating at 95 °C for 2 min. 1 μL of the resulting mixture was further diluted to give 0.005 μM 5' $^{32}$P-labeled oligonucleotides. Samples were resolved by urea-PAGE and analysed by autoradiographic imaging.

**Nucleotide Screening in RNA Ligation Assay (Fig. 2b)**

In the nucleotide screening study, 0.1 μM 5' $^{32}P$-labeled RNA oligo1 and 0.5 μM RNA oligo2 were incubated with 1.0 μM C12orf29$^{WT}$ and 200 μM of the indicated nucleotides in 1x RNA ligation buffer at 37 °C for 1 h in a total volume of 10 μL. The reaction was stopped by adding 150 μL stopping solution and heating at 95 °C for 2 min. 1 μL of samples were resolved by urea-PAGE and analysed by autoradiographic imaging.

**Autoradiographic Analysis**

*Urea-PAGE Analysis and Phosphorimaging*

Samples were quenched by adding stopping solution and denaturing at 95 °C for 2 min. 1 μL of denatured sample were loaded onto a pre-warmed 12% urea-PAGE sequencing gel. Electrophoresis was performed in 1x TBE buffer (90 mM Tris-HCl pH 8.0, 90 mM boric acid, 2.0 mM EDTA) at constant power of 100 W. The gel was transferred onto Whatman® filter paper and dried at 80 °C for 2 h under vacuum. After cooling down to room temperature, the gel was exposed to a storage phosphor screen, of which the readout was performed by Typhoon™ FLA-9500 in phosphorimaging mode and further processed by Image Lab, Bio-Rad.

*SDS-PAGE and Phosphorimaging*

Samples were mixed with 6x loading buffer to a final concentration of 1x loading buffer and denatured at 95 °C for 5 min. Separation of proteins were realized by using 4% acrylamide stacking gel and 12.5% acrylamide resolving gel. As references an unstained or pre-stained protein ladder was used. Electrophoresis was performed in 1x SDS running buffer at constant voltage of 220 V for 40 min. The gel was stained by Coomassie staining solution at room temperature for 30 min after electrophoresis, followed by destaining, rinsing with MilliQ® $H_2O$, and imaging. In turn, the gel was soaked in the fixing solution (3% (v/v) glycerol, 20% (v/v) HOAc, 20% (v/v) MeOH) at room temperature for 30 min and rinsed in MilliQ® $H_2O$ for 5 min before being transferred onto Whatman® filter paper and drying at 80 °C for 1 h under vacuum. After cooling down to room temperature, the gel was exposed to a storage phosphor screen, of which the readout was performed by Typhoon™ FLA-9500 in phosphorimaging mode and further processed by Image Lab, Bio-Rad.

**Mutational Effects on RNA Ligation**

0.1 µM 5' $^{32}$P-labeled RNA14 were applied in the standard RNA ligation assay with 1.0 µM C12orf29 variants. The readout was processed by Image Lab, Bio-Rad to give signal intensities, which were normalized to that of control experiment (without enzyme). The values represented arithmetic mean ± SD from three biological replicates in histogram.

##### Preparation of *C12ORF29* Knockout HEK293 Cell Lines

The *C12ORF29* knockout cell lines were generated in collaboration with trenzyme GmbH (Konstanz, Germany): gRNA oligonucleotides were designed to target and knock out *C12ORF29*. After successful nucleofection of the gRNA into the cells, single cell cloning by limiting dilution was performed. All resulting clones were picked and analysed resulting in clones that were homozygous for *C12ORF29* knockout. One of the knockout cell lines was chosen for further experiments.

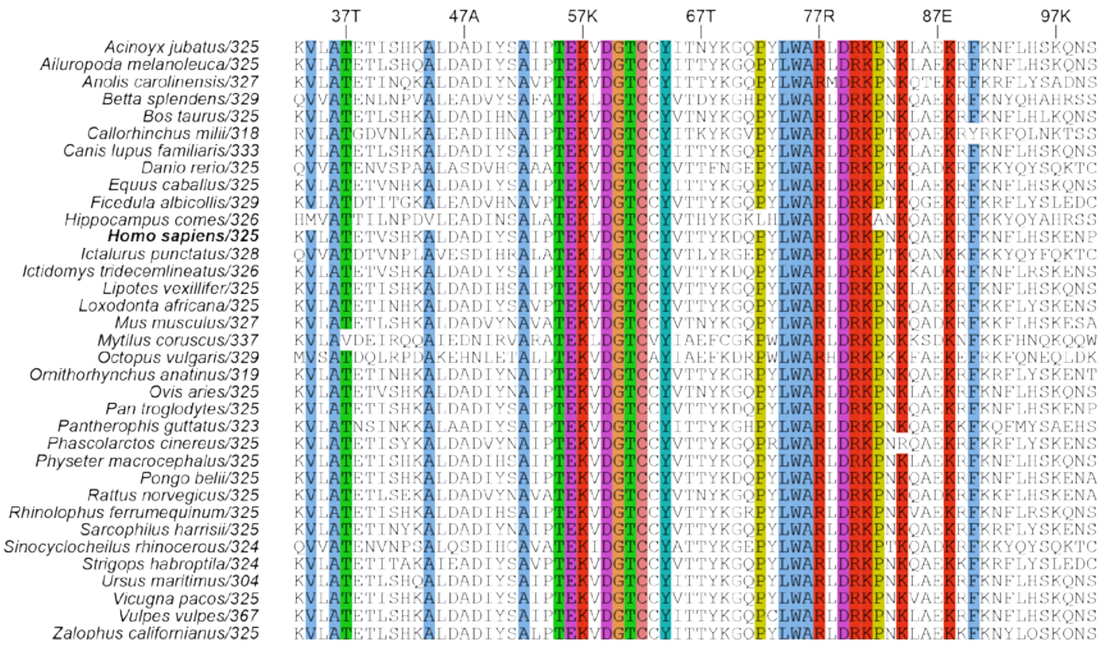

**Supplementary Fig. 1**

**Multiple amino acid sequence alignment of C12orf29 homologues in different species.** The amino acid sequence of C12orf29 of *Homo sapiens* (in bold) was used as reference. The number after the species names represents the length of C12orf29 homologues. Residues with >95% identities are shaded with different colors. Hydrophobic residues are in blue. Positively charged residues are in red. Negatively charged residues are in magenta. Polar residues are in green. Cysteines are in pink. Glycines are in orange. Prolines are in yellow. Aromatic residues are in cyan. The alignment was done with Clustal Omega[29].

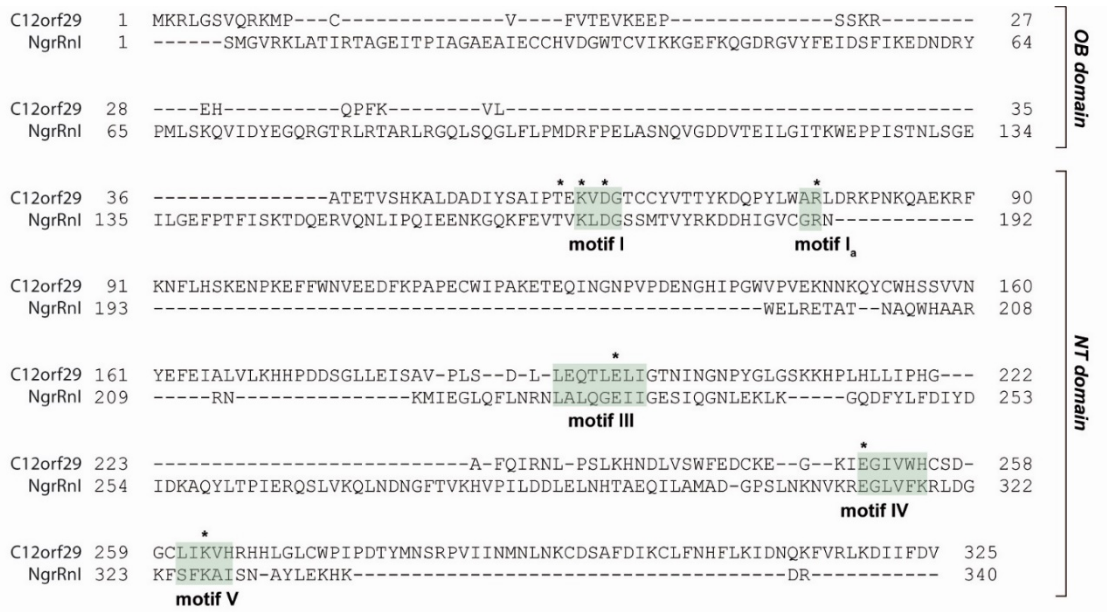

**Supplementary Fig. 2**

**Structure-based amino acid sequence alignment of C12orf29 and *Ngr*Rnl**. The OB domain and NT domain of *Ngr*Rnl are indicated by brackets at right. Motifs I, $I_a$, III, IV, and V are shaded green. Residues that are potentially essential for the ligase activity were indicated by asterisks. The alignment was done with PROMALS3D[30] using the C12orf29 model predicted by AlphaFold and *Ngr*Rnl with PDB ID: 5COT. OB, oligonucleotide-binding. NT, nucleotidyltransferase.

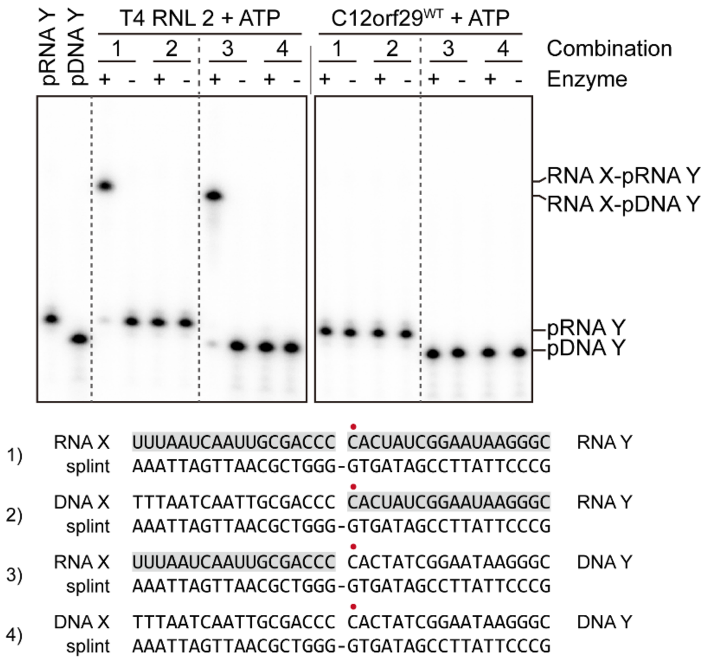

**Supplementary Fig. 3**

**Non-reactive substrates for ligation.** Four nicked duplexes with different combinations of DNA/RNA oligonucleotides were tested separately. RNA oligos were shaded in gray. The $^{32}$P-labeled 5'-$PO_4$ were depicted in red. All reactions were performed under identical conditions: 1.0 μM C12orf29$^{WT}$, 0.1 μM 5' $^{32}$P-labeled oligonucleotide, 200 μM ATP, 5.0 mM $MgCl_2$, and 1.0 mM DTT in 50 mM Tris-HOAc at pH 7.0 for 60 min at 37 $^{o}$C. T4 RNA ligase 2 was employed for the positive controls. The radioactive oligonucleotides were resolved by urea-PAGE and phosphorimaging.

| Oligonucleotide | Sequence (5' to 3') |
|---|---|
| **RNA 1** | GGC ACU CAG ACU CAG AG |
| **RNA 2** | CUC UGA GUA A |
| **RNA 3** | AGC ACU CAG ACU CAG ACU AGU AA |
| **RNA 4** | UUA CUA GUC UGA GUA A |
| **RNA 5** | AGC ACU CAG ACU AGU GAG AAC UAG UAA |
| **RNA 6** | CGC ACU CAG ACU AGU GAG AAC UAG UAA |
| **RNA 7** | GGC ACU CAG ACU AGU GAG AAC UAG UAA |
| **RNA 8** | UGC ACU CAG ACU AGU GAG AAC UAG UAA |
| **RNA 9** | AGC ACU CAG ACU AGU GAG AAC UAG UAC |
| **RNA 10** | AGC ACU CAG ACU AGU GAG AAC UAG UAG |
| **RNA 11** | AGC ACU CAG ACU AGU GAG AAC UAG UAU |
| **RNA 12** | AGC CAG ACU AGU GAG AAC UAG UAA |
| **RNA 13** | AGC ACU AGU GAG AAC UAG UAA |
| **RNA 14** | AGC ACU CAG ACU AGU AAA GCA CUC AGA CUA GUA A |
| **DNA 1** | GGC ACT CAG ACT CAG AG |
| **DNA 2** | CTC TGA GTA A |

**Supplementary Fig. 4**

**Oligonucleotide sequences used in this study.**

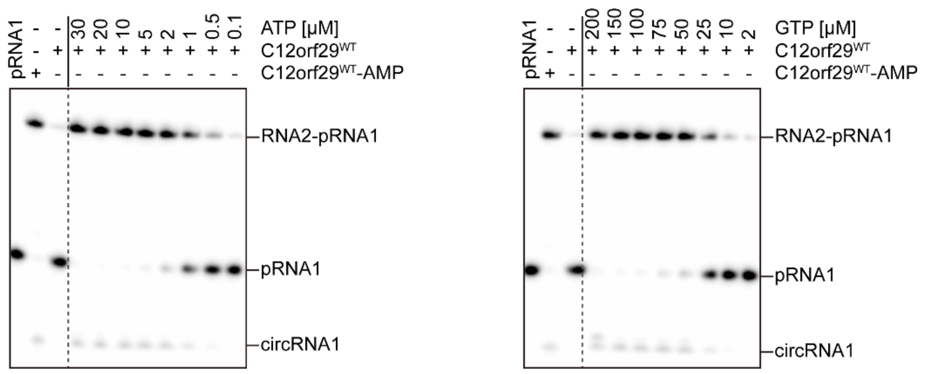

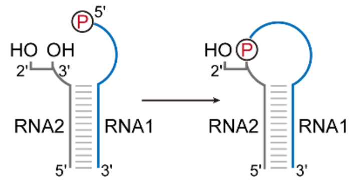

| | ATP | GTP |
|---|---|---|
| $V_{max}$ [nM/min] | 1.68 ± 0.06 | 1.89 ± 0.11 |
| $K_M$ [μM] | 1.30 ± 0.19 | 37.7 ± 7.26 |
| $k_{cat}$ [$s^{-1}$] | $2.81 \times 10^{-5}$ ± $9.88 \times 10^{-7}$ | $3.14 \times 10^{-5}$ ± $1.90 \times 10^{-6}$ |
| $k_{cat}/K_M$ [$s^{-1}$/M] | 21.6 | 0.83 |

**Supplementary Fig. 5**

**Steady state kinetics using ATP or GTP as substrate.** Non-cropped urea-PAGE analysis depicted in Fig. 2c were displayed in the top panel. Exemplary RNA ligation reactions depicted were performed with 1.0 μM C12orf29$^{WT}$ for 60 min at 37 °C at varying concentrations of ATP or GTP. The rate of RNA2-pRNA1 formation was quantified by the intensities of the bands for each chamber. Values represented arithmetic mean ± SD from biological triplicates.

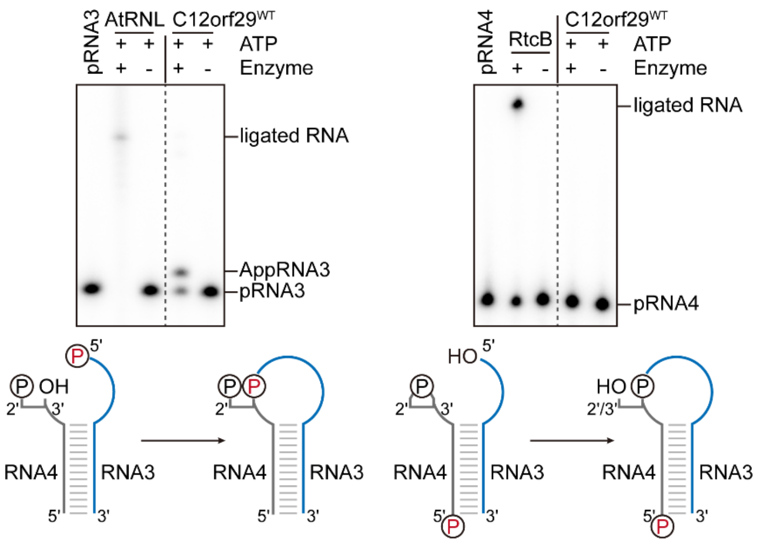

**Supplementary Fig. 6**

**Substrate scope of C12orf29.** Neither RNA termini modifications (2'-$PO_4$ or 2',3'-c$PO_4$) were not used as substrates for C12orf29. Radioactive 5'-$PO_4$ are shown in red.

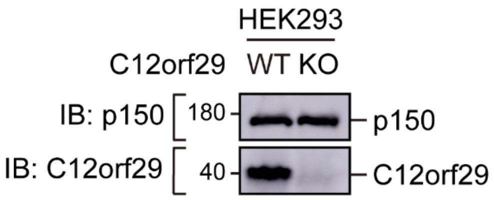

**Supplementary Fig. 7**

**Western blot analysis of C12orf29 expression in WT and *C12ORF29*-KO HEK293 cells.** WT and *C12ORF29*-KO HEK293 cells were lysed in 1x loading buffer without β-mercaptoethanol, heated to 95 °C for 5 min and sonicated on ice. The lysates were centrifuged (13,400 rpm, 5 min) and the protein concentrations of the supernatant were determined by BCA assay. 40 µg total protein were resolved by SDS-PAGE and subsequent Western Blotting as described above. p150 was used as loading control.

| Sample | WT 0 µM | KO 0 µM | WT 10 µM | KO 10 µM | WT 20 µM | KO 20 µM | WT 40 µM | KO 40 µM | WT 60 µM | KO 60 µM | WT 100 µM | KO 100 µM |
|---|---|---|---|---|---|---|---|---|---|---|---|---|
| Sample conc. [pg/µl] | 5200 | 4360 | 6120 | 4500 | 3910 | 4750 | 4680 | 4460 | 4470 | 4000 | 4800 | 4240 |
| 18S [pg/µl] | 841 | 724 | 905 | 680 | 614 | 802 | 727 | 968 | 799 | 884 | 938 | 1120 |
| 28S [pg/µl] | 3040 | 2210 | 3500 | 2260 | 1950 | 2050 | 2220 | 984 | 1970 | 900 | 745 | 540 |
| 18S/total RNA | 0,16 | 0,17 | 0,15 | 0,15 | 0,16 | 0,17 | 0,16 | 0,22 | 0,18 | 0,22 | 0,20 | 0,26 |
| 28S/total RNA | 0,58 | 0,51 | 0,57 | 0,50 | 0,50 | 0,43 | 0,47 | 0,22 | 0,44 | 0,23 | 0,16 | 0,13 |
| 28S/18S | 3,60 | 3,00 | 3,90 | 3,30 | 3,20 | 2,60 | 3,10 | 1,00 | 2,50 | 1,00 | 0,80 | 0,50 |

#### Supplementary Fig. 8

**RNA integrity analysis of total RNA of WT and KO HEK293 cells after menadione treatment**. Data from the experiments in Fig. 4c was shown in Supplementary Fig. 8, indicating the total concentrations of RNA, the concentrations of 18S and 28S rRNA, the ratios of 18S and 28S rRNA to the total RNA and the 28S/18S ratio. The RNA concentrations, as well as the 28S/18S ratio, were directly calculated by the TapeStation system. The ratios of 18S and 28S to total RNA were calculated respectively from the shown concentrations.

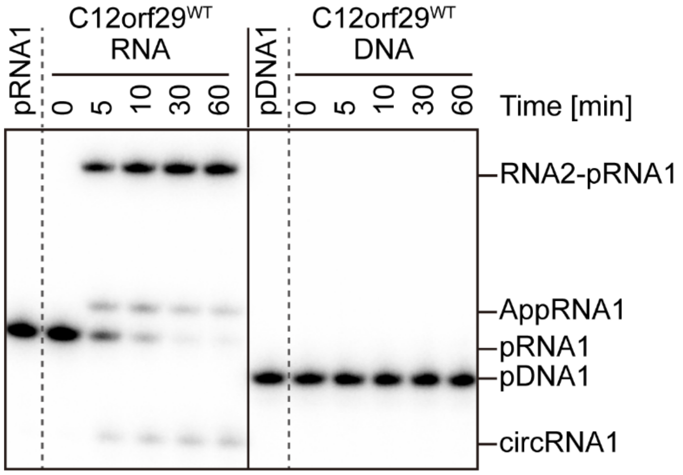

**Supplementary Fig. 9**

**Non-cropped urea-PAGE analysis depicted in Fig. 2a.**

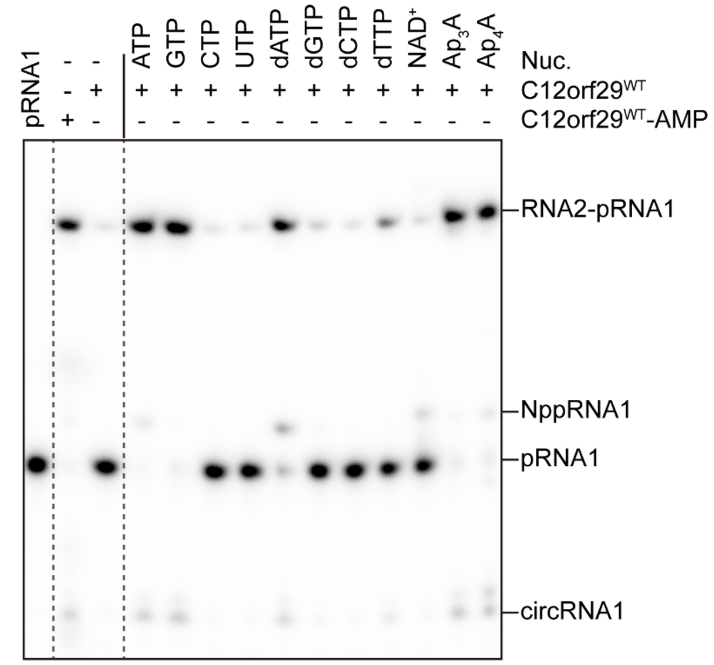

**Supplementary Fig. 10**

**Non-cropped urea-PAGE analysis depicted in Fig. 2b.**

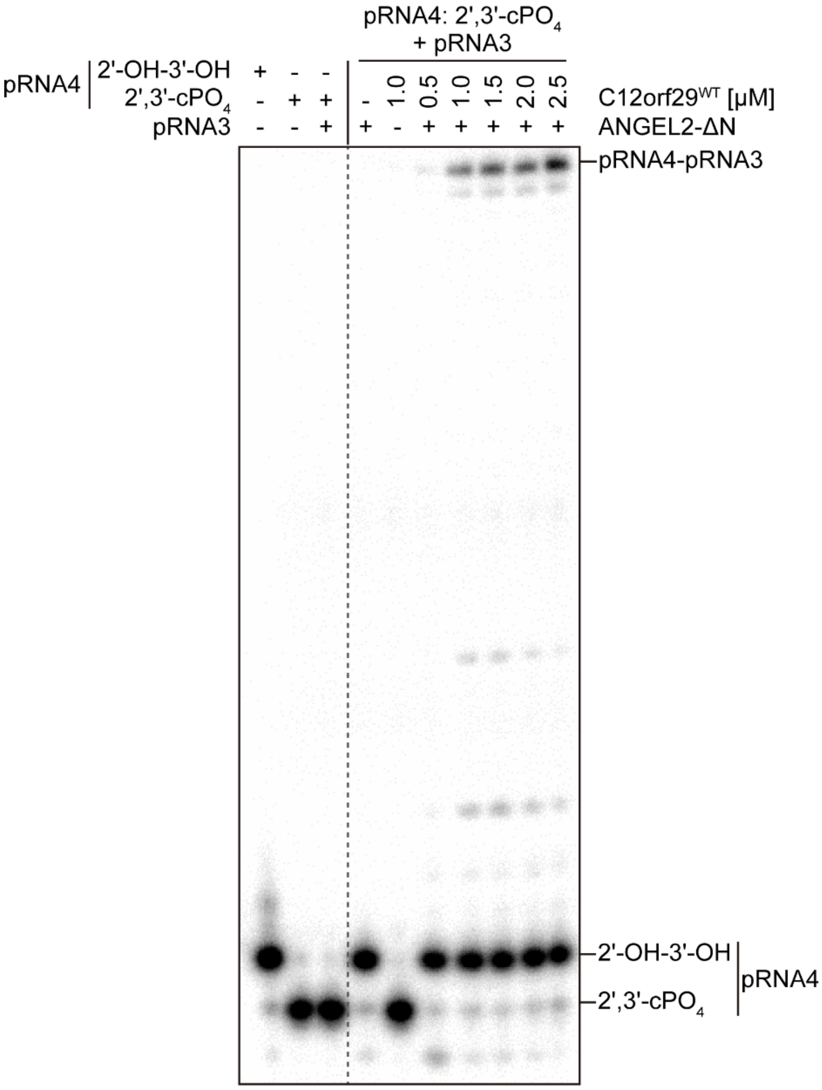

**Supplementary Fig. 11**

**Non-cropped urea-PAGE analysis depicted in Fig. 2d.**

#### Synthesis and analysis of chemical probes

#### Chemicals and Reagents

Unless otherwise noted, all chemicals were purchased from commercial suppliers (abcr, Acros Organics, Fluka, Carl Roth, Sigma-Aldrich, TCI, and VWR) and used without further purification. Dry solvents were used in all reactions unless otherwise noted. DIPEA was dried over $CaH_2$, distilled, and stored over molecular sieves (4 Å) under $N_2$. $Et_3N$ was dried over KOH, distilled, and stored over molecular sieves (4 Å) under $N_2$. The $CDCl_3$ and DMSO-$d_6$ were stored over molecular sieves (4 Å) under $N_2$ if applied in the NMR measurement of $P^{III}$ compounds. For the preparation of aqueous solutions and buffers, ultra-pure water generated with the Merck Millipore BioPak® system (MilliQ® $H_2O$) was used.

#### General Remarks

NMR spectra were recorded by Bruker Avance III 400 MHz, 500 MHz, and 600 MHz spectrometers at room temperature. Chemical shifts (in ppm) were calibrated using residual undertreated solvent in $CDCl_3$, DMSO-$d_6$, and $D_2O$ as references, respectively. Multiplicities are denoted as s = singlet, d = doublet, t = triplet, q = quartet, quint = quintet, m = multiplet, dd = doublet of doublets, dt = doublet of triplets, ddd = doublet of doublet of doublets, br = broad. Low resolution MS (LR-MS) was performed by amazon SL (ESI-ion trap) from Bruker Daltonics. High resolution MS (HR-MS) were performed by micrOTOF II (ESI-TOF) from Bruker Daltonics.

#### Synthesis of modified $Ap_3A$

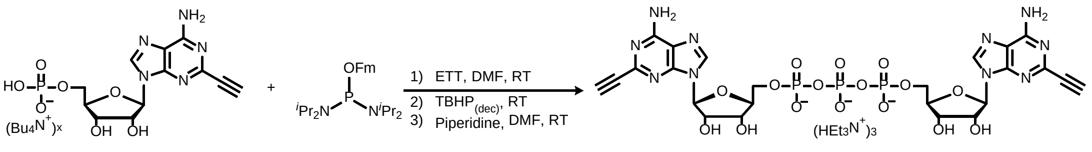

1 2 *C*2-eAp$_3$A

**Supplementary Scheme 1 Synthesis of *C*2-eAp3A.**

Compound **1**[31] as tetrabutylammonium salt (2.0 equiv, 0.050 mmol, 36.5 mg) was co-evaporated with MeCN (3 × 3 mL) in an oven-dried 5 mL flask and dried under vacuum for 1 h. The compound was then dissolved in 0.5 mL DMF under $N_2$, followed by the addition of ETT (2.7 equiv, 67.5 μmol, 8.8 mg) and **2**[32] (1.0 equiv, 0.025 mmol, 10.7 mg). The resulting mixture was stirred at room temperature for 20 min, and $TBHP_{(dec)}$ (15.0 equiv, 0.375 mmol, 68.1 μL) was added in three portions. After stirring for 30 min, the reaction mixture was transferred to a falcon tube before the addition of $Et_2O$/*n*-

hexane (5:1). The resulting suspension was centrifuged to give a pellet, which was washed with $Et_2O$/*n*-hexane (5:1) and centrifuged three times. After removing the supernatant, the pellet was dissolved in 0.5 mL DMF, and 50 μL piperidine was added to the solution. The reaction was stirred for 10 min at room temperature and was quenched by the addition of $Et_2O$/*n*-hexane (5:1). The suspension was centrifuged to give a pellet, which was washed with $Et_2O$/*n*-hexane (5:1) and centrifuged three times. The resulting pellet was purified via IEX-FPLC and RP-HPLC to give 9.7 mg **5** in 20% yield (UV absorption estimated). **$^1$H NMR** (400 MHz, $D_2O$) δ 8.38 (s, 2H, H-8, H-8''), 5.98 (d, *J* = 4.4 Hz, 2H, H-1', H-1'''), 4.63 (t, *J* = 4.6 Hz, 2H, H-2', H-2'''), 4.54 (t, *J* = 5.0 Hz, 2H, H-3', H-3'''), 4.41-4.34 (m, 6H, H-4', H-4''', $H_a$-5', $H_a$-5''', $H_b$-5', $H_b$-5'''), 3.59 (s, 2H, *C*2-C≡CH, *C*2''-C≡CH). **$^{13}$C NMR** (100 MHz, $D_2O$) δ 154.8 (C-6, C-6''), 148.3 (C-4, C-4''), 144.9 (C-2, C-2''), 140.2 (C-8, C-8''), 117.9 (C-5, C-5''), 87.4 (C-1', C-1'''), 83.2 (d, *J* = 9.0 Hz, C-4', C-4'''), 80.8 (*C*2-*C*≡CH, *C*2''-*C*≡CH), 76.3 (*C*2-C≡*C*H, *C*2''-C≡*C*H), 75.1 (C-2', C-2'''), 69.8 (C-3', C-3'''), 64.8 (d, *J* = 5.0 Hz, C-5', C-5'''). **$^{31}$P NMR** (162 MHz, $D_2O$) δ -11.52 (d, *J* = 19.4 Hz, 2P, P-α, P-α'), -23.04 (t, *J* = 19.4 Hz, 1P, P-β). **HR-MS** *m*/*z* calcd for $C_{24}H_{26}N_{10}O_{16}P_3^-$ $(M - H)^-$ 803.0747, found 803.0742, deviation 0.6 ppm.

#### Synthesis of modified Azide

The synthesis was inspired by ref. 33

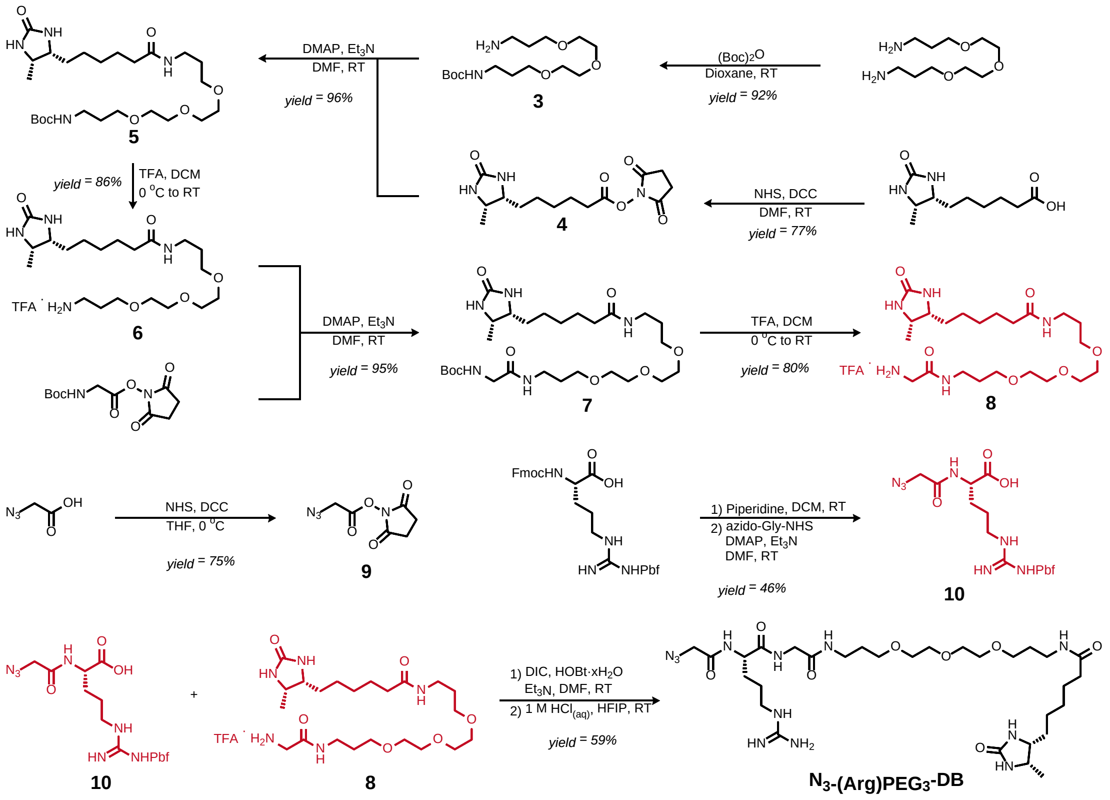

**Supplementary Scheme 2 Overview of the synthesis of $N_3$-(Arg)PEG$_3$-DB.**

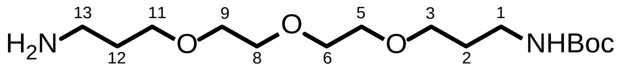

*tert*-Butyl (3-(2-(2-(3-aminopropoxy)ethoxy)ethoxy)propyl)carbamate **3**[34]

An oven-dried 250 mL two-neck flask was evacuated and flushed with $N_2$ trice, followed by the addition of 4,7,10-trioxa-1,13-tridecanediamine (5.2 equiv, 52.0 mmol, 11.38 mL) and 50 mL dioxane. A solution of di-*tert*-butyl dicarbonate (1.0 equiv, 10.0 mmol, 2.18 g) in 20 mL dioxane was added dropwise within 3 h at room temperature. The reaction mixture was stirred for additional 5 h and then concentrated. The residue was dissolved in 50 mL $H_2O$ and extracted with DCM (4 × 50 mL). The organic phases were collected and washed with brine (4 × 30 mL). The extraction and the subsequent washing procedure were repeated four times. The organic phases were combined, dried over $MgSO_4$, concentrated and purified via silica chromatography (DCM:MeOH = 1:1) to give 2.94 g **3** as yellow oil in 92% yield. **$^1$H NMR** (400 MHz, $CDCl_3$) δ 5.11 (br s, 1H, NH-1), 3.63-3.60 (m, 4H, H-6, H-8), 3.58-3.56 (m, 4H, H-5, H-9), 3.54-3.50 (m,

4H, H-3, H-11), 3.22-3.17 (m, 2H, H-1), 2.81 (t, $J$ = 6.6 Hz, 2H, H-13), 2.16 (br s, 2H, $NH_2$-13), 1.76-1.70 (m, 4H, H-2, H-12), 1.41 (s, 9H, -C($CH_3$)$_3$). **$^{13}$C NMR** (100 MHz, $CDCl_3$) δ 156.2 (-<u>C</u>OO-), 79.0 (-<u>C</u>($CH_3$)$_3$), 70.68, 70.65 (C-6, C-8), 70.31 (C-5), 70.26 (C-9), 69.64, 69.61 (C-3, C-11), 39.7 (C-13), 38.6 (C-1), 32.8 (C-12), 29.8 (C-2), 28.6 (-C(<u>C</u>$H_3$)$_3$). **LR-MS** *m*/*z* calcd for $C_{15}H_{33}N_2O_5^+$ $(M + H)^+$ 321.24, found 321.21.

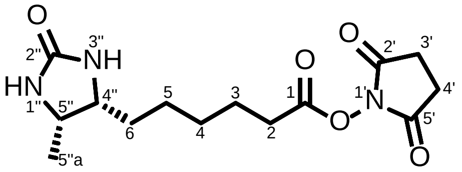

2,5-Dioxopyrrolidin-1-yl 6-((4*R*,5*S*)-5-methyl-2-oxoimidazolidin-4-yl)hexanoate **4**[35]

Desthiobiotin (1.0 equiv, 3.55 mmol, 761.2 mg), *N*-hydroxysuccinimide (1.0 equiv, 3.55 mmol, 408.7mg), and DCC (1.3 equiv, 4.62 mmol, 952.9 mg) were added to an oven-dried 50 mL flask, which was evacuated and flushed with $N_2$ trice, followed by the addition of 30 mL DMF. After stirring for 48 h at room temperature, the reaction mixture was filtered to remove formed *N*,*N*'-dicyclohexylurea. The solvent was evaporated, and 125 mL $Et_2O$ was added. The suspension was filtered after stirring for 2 h at room temperature. The resulting white solid was recrystallized in 50 mL $^i$PrOH to give 851.3 mg **4** as white powder in 77% yield. **$^1$H NMR** (400 MHz, DMSO-$d_6$) δ 6.29 (s, 1H, NH-3"), 6.11 (s, 1H, NH-1"), 3.61 (quint, $J$ = 6.7 Hz, 1H, H-5"), 3.51-3.46 (m, 1H, H-4"), 2.81 (s, 4H, H-3', H-4'), 2.66 (t, $J$ = 7.4 Hz, 2H, H-2), 1.62 (quint, $J$ = 7.1 Hz, 2H, H-3), 1.42-1.18 (m, 6H, H-4, H-5, H-6), 0.96 (d, $J$ = 6.0 Hz, 3H, H-5"a). **$^{13}$C NMR** (100 MHz, DMSO-$d_6$) δ 170.2 (C-2', C-5'), 169.0 (C-1), 162.7 (C-2"), 54.9 (C-4"), 50.2 (C-5"), 30.1 (C-2), 29.4 (C-6), 28.0 (C-4), 25.4 (C-3', C-4'), 25.3 (C-5), 24.2 (C-3), 15.5 (C-5"a). **HR-MS** *m*/*z* calcd for $C_{14}H_{22}N_3O_5^+$ $(M + H)^+$ 312.1559, found 312.1553, deviation 1.9 ppm.

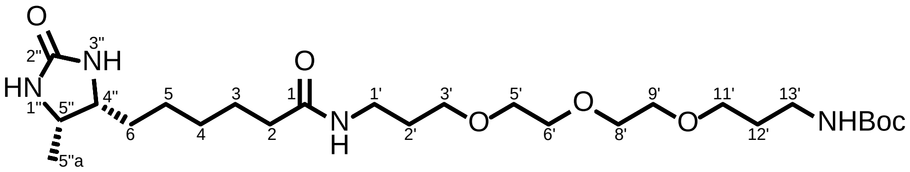

*tert*-Butyl (20-((4*R*,5*S*)-5-methyl-2-oxoimidazolidin-4-yl)-15-oxo-4,7,10-trioxa-14-azaicosyl)carbamate **5**[35]

Compound **4** (1.0 equiv, 3.19 mmol, 991.2 mg) and DMAP (0.1 equiv, 0.32 mmol, 39.0 mg) were added to an oven-dried 50 mL flask, which was evacuated and flushed with $N_2$ trice, followed by the addition of a solution of compound **3** (2.0 equiv, 6.38 mmol, 2.04 g) in 10 mL DMF and $Et_3N$ (1.0 equiv, 3.19 mmol, 443.4 μL). After stirring overnight at room temperature, DMF was removed, and the residue was dissolved in 50 mL DCM. The solution was successively washed with 0.5 M HCl (2 × 25 mL), 1 M $K_2CO_3$ (2 × 25 mL), and brine (2 × 25 mL). The resulting organic phase was dried over $MgSO_4$, concentrated, and purified via silica chromatography (DCM:MeOH = 20:1) to give 1.58 g **5** as pale yellow oil in 96% yield. **$^1$H NMR** (400 MHz, DMSO-*d*$_6$) δ 7.70 (t, *J* = 5.6 Hz, 1H, NH-1'), 6.73 (t, *J* = 5.8 Hz, 1H, NH-13'), 6.28 (s, 1H, NH-3''), 6.10 (s, 1H, NH-1''), 3.60 (quint, *J* = 6.7 Hz, 1H, H-5''), 3.52-3.44 (m, 9H, H-5', H-6', H-8', H-9', H-4''), 3.40-3.36 (m, 4H, H-3', H-11'), 3.06 (q, *J* = 6.5 Hz, 2H, H-1'), 2.95 (q, *J* = 6.5 Hz, 2H, H-13'), 2.03 (t, *J* = 7.4 Hz, 2H, H-2), 1.63-1.55 (m, 4H, H-2', H-12'), 1.52-1.44 (m, 2H, H-3), 1.37 (s, 9H, -C($CH_3$)$_3$), 1.35-1.16 (m, 6H, H-4, H-5, H-6), 0.95 (d, *J* = 6.4 Hz, 3H, H-5''a). **$^{13}$C NMR** (100 MHz, DMSO-*d*$_6$) δ 171.9 (C-1), 162.8 (C-2''), 155.5 (-<u>C</u>OO-), 77.4 (-<u>C</u>($CH_3$)$_3$), 69.7 (C-6', C-8'), 69.5 (C-5', C-9'), 68.09, 68.06 (C-3', C-11'), 55.0 (C-4''), 50.2 (C-5''), 37.2 (C-13'), 35.7 (C-1'), 35.3 (C-2), 29.7 (C-12'), 29.5 (C-6), 29.4 (C-2'), 28.7 (C-4), 28.2 (-C(<u>C</u>$H_3$)$_3$), 25.5 (C-5), 25.2 (C-3), 15.5 (C-5''a). **HR-MS** *m/z* calcd for $C_{25}H_{49}N_4O_7^+$ $(M + H)^+$ 517.3601, found 517.3605, deviation 0.8 ppm.

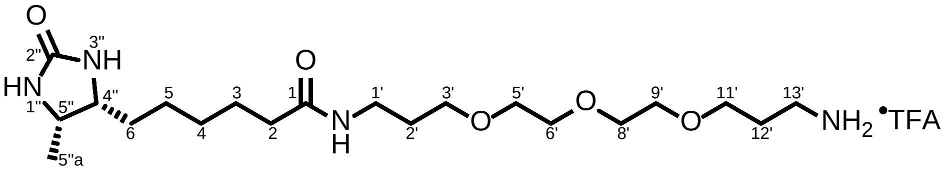

*N*-(3-(2-(2-(3-Aminopropoxy)ethoxy)ethoxy)propyl)-6-((4*R*,5*S*)-5-methyl-2-oxoimidazolidin-4-yl)hexanamide (trifluoroacetic acid salt form) **6**[35]

Compound **5** (1.0 equiv, 3.05 mmol, 1.57 g) was added to an oven-dried 50 mL flask, followed by the addition of 25 mL DCM. To the solution, 12.5 mL TFA was added slowly at 0 °C. The reaction mixture was warmed up to room temperature and stirred for 2 h. Then DCM and TFA was removed under high vacuum overnight. The residue was purified via silica chromatography ($CHCl_3$:MeOH:$NH_{3(aq)}$ = 1:0.25:0.1) to give 1.40 g **6** as sticky yellow oil in 86% yield. **$^1$H NMR** (400 MHz, DMSO-*d*$_6$) δ 7.75 (t, *J* = 5.6 Hz, 1H, NH-1'), 6.29 (s, 1H, NH-3''), 6.12 (s, 1H, NH-1''), 4.67 (br s, 3H, $NH_3^+$-13'), 3.60 (quint, *J* = 6.7 Hz, 1H, H-5''), 3.53-3.44 (m, 11H, H-5', H-6', H-8', H-9', H-11', H-4''), 3.38 (t, *J* = 6.4 Hz, 2H, H-3'), 3.06 (q, *J* = 6.4 Hz, 2H, H-1'), 2.77 (t, *J* = 7.2 Hz, 2H, H-13'), 2.03 (t, *J* = 7.4 Hz, 2H, H-2), 1.71 (quint, *J* = 6.6 Hz, 2H, H-12'), 1.60 (quint, *J* = 6.7 Hz, 2H, H-2'), 1.47 (quint, *J* = 7.3 Hz, 2H, H-3), 1.36-1.16 (m, 6H, H-4, H-5, H-6), 0.95 (d, *J* = 6.4 Hz, 3H, H-5''a). **$^{13}$C NMR** (100 MHz, DMSO-*d*$_6$) δ 172.0 (C-1), 162.8 (C-2''), 158.0 (q, $^3J_{FC}$ = 30.7 Hz, -<u>C</u>O$CF_3$), 117.3 (q, $^2J_{FC}$ = 298.8 Hz, -<u>C</u>$F_3$), 69.74, 69.68 (C-6', C-8'), 69.52 (C-5'), 69.47 (C-9'), 68.1 (C-3'), 67.7 (C-11'), 55.0 (C-4''), 50.2 (C-5''), 37.4 (C-13'), 35.7 (C-1'), 35.3 (C-2), 29.5 (C-6), 29.4 (C-2'), 29.0

(C-12'), 28.7 (C-4), 25.6 (C-5), 25.2 (C-3), 15.5 (C-5"a). **$^{19}$F NMR** (376 MHz, DMSO-*d*$_6$) δ -73.6 (-$CF_3$). **HR-MS** *m*/*z* calcd for $C_{20}H_{41}N_4O_5^+$ (M + H)$^+$ 417.3077, found 417.3069, deviation 1.9 ppm.

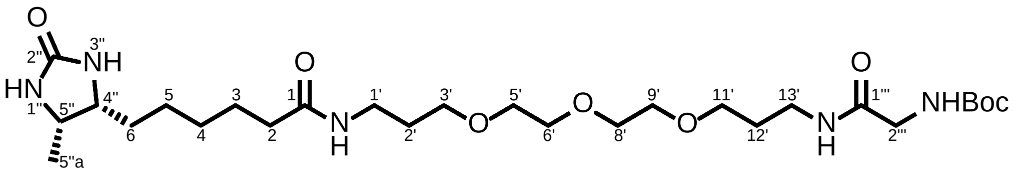

*tert*-Butyl (23-((4*R*,5*S*)-5-methyl-2-oxoimidazolidin-4-yl)-2,18-dioxo-7,10,13-trioxa-3,17-diazatricosyl)carbamate **7**

DMAP (0.1 equiv, 0.13 mmol, 15.9 mg) and Boc-Gly-NHS (1.5 equiv, 1.95 mmol, 530.6 mg) was added to an oven-dried 25 mL flask, which was evacuated and flushed with $N_2$ trice, followed by the addition of a solution of compound **6** (1.0 equiv, 1.30 mmol, 689.4 mg) in 10 mL DMF and $Et_3N$ (2.5 equiv, 3.25 mmol, 451.7 μL). After stirring overnight at room temperature, DMF was removed, and the residue was dissolved in 50 mL DCM. The solution was washed with brine (1 × 25 mL), which was extracted with DCM (4 × 50 mL). The combined organic phases were dried over $MgSO_4$, concentrated, and purified via silica chromatography (DCM:MeOH = 20:1) to give 707.6 mg **7** as sticky colorless oil in 95% yield. **$^1$H NMR** (400 MHz, DMSO-*d*$_6$) δ 7.72-7.70 (m, 2H, NH-1', NH-13'), 6.87 (t, *J* = 6.2 Hz, 1H, NH-2'''), 6.28 (s, 1H, NH-3"), 6.10 (s, 1H, NH-1"), 3.60 (quint, *J* = 6.8 Hz, 1H, H-5"), 3.53-3.45 (m, 11H, H-5', H-6', H-8', H-9', H-4", H-2'''), 3.40-3.36 (m, 4H, H-3', H-11'), 3.12-3.04 (m, 4H, H-1', H-13'), 2.03 (t, *J* = 7.4 Hz, 2H, H-2), 1.65-1.57 (m, 4H, H-2', H-12'), 1.47 (quint, *J* = 7.3 Hz, 2H, H-3), 1.38 (s, 9H, -$C(CH_3)_3$), 1.36-1.16 (m, 6H, H-4, H-5, H-6), 0.95 (d, *J* = 6.4 Hz, 3H, H-5"a). **$^{13}$C NMR** (100 MHz, DMSO-*d*$_6$) δ 171.9 (C-1), 169.1 (C-1'''), 162.8 (C-2"), 155.5 (-$\underline{C}$OO-), 78.0 (-$\underline{C}(CH_3)_3$), 69.74, 69.73 (C-6', C-8'), 69.5 (C-5', C-9'), 68.1 (C-3', C-11'), 55.0 (C-4"), 50.2 (C-5"), 43.3 (C-2'''), 35.9 (C-13'), 35.7 (C-1'), 35.3 (C-2), 29.5 (C-6), 29.4 (C-2'), 29.3 (C-12'), 28.7 (C-4), 28.2 (-$C(\underline{C}H_3)_3$), 25.6 (C-5), 25.2 (C-3), 15.5 (C-5"a). **HR-MS** *m*/*z* calcd for $C_{27}H_{52}N_5O_8^+$ (M + H)$^+$ 574.3810, found 574.3821, deviation 2.0 ppm.

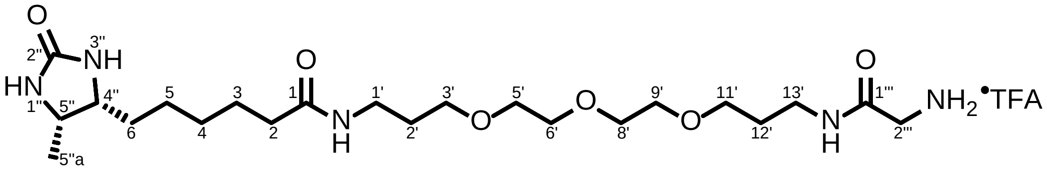

*N*-(1-Amino-2-oxo-7,10,13-trioxa-3-azahexadecan-16-yl)-6-((4*R*,5*S*)-5-methyl-2-oxoimidazolidin-4-yl)hexanamide (trifluoroacetic acid salt form) **8**

Compound **7** (1.0 equiv, 1.22 mmol, 697.0 mg) was added to an oven-dried 50 mL flask, followed by the addition of 10 mL DCM. To the solution, 5.0 mL TFA was added slowly at 0 °C. The reaction mixture was warmed up to room temperature and stirred for 2 h. Then DCM and TFA was removed under high vacuum overnight. The residue was purified via silica chromatography ($CHCl_3$:MeOH:$NH_{3(aq)}$ = 1:0.25:0.1) to give 568.3 mg of **8** sticky pale yellow oil in 80% yield. **$^1$H NMR** (400 MHz, DMSO-*d*$_6$) δ 7.86 (t, *J* = 5.8 Hz, 1H, NH-1'), 7.72 (t, *J* = 5.6 Hz, 1H, NH-13'), 6.28 (s, 1H, NH-3''), 6.10 (s, 1H, NH-1''), 3.60 (quint, *J* = 6.7 Hz, 1H, H-5''), 3.53-3.45 (m, 9H, H-5', H-6', H-8', H-9', H-4''), 3.42-3.36 (m, 4H, H-3', H-11'), 3.28 (br s, 3H, $NH_3^+$-2'''), 3.13 (q, *J* = 6.4 Hz, 2H, H-1'), 3.10 (s, 2H, H-2'''), 3.06 (q, *J* = 6.4 Hz, 2H, H-13'), 2.03 (t, *J* = 7.4 Hz, 2H, H-2), 1.67-1.57 (m, 4H, H-2', H-12'), 1.51-1.44 (m, 2H, H-3), 1.36-1.15 (m, 6H, H-4, H-5, H-6), 0.95 (d, *J* = 6.4 Hz, 3H, H-5''a). **$^{13}$C NMR** (100 MHz, DMSO-*d*$_6$) δ 171.93 (C-1), 171.85 (C-1'''), 162.8 (C-2''), 69.8 (C-6', C-8'), 69.53 (C-5', C-9'), 68.2 (C-3'), 68.1 (C-11'), 55.0 (C-4''), 50.2 (C-5''), 44.2 (C-2'''), 35.8 (C-13'), 35.7 (C-1'), 35.3 (C-2), 29.5 (C-6), 29.4 (C-2'), 29.3 (C-12'), 28.7 (C-4), 25.6 (C-5), 25.2 (C-3), 15.5 (C-5''a). $^{19}$F NMR (376 MHz, DMSO-*d*$_6$) δ -73.5 (-$CF_3$). **HR-MS** *m*/*z* calcd for $C_{22}H_{44}N_5O_6^+$ $(M + H)^+$ 474.3286, found 474.3282, deviation 0.8 ppm.

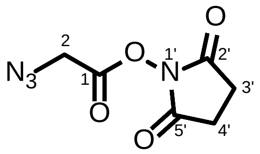

2,5-Dioxopyrrolidin-1-yl 2-azidoacetate **9**[36]

*N*-Hydroxysuccinimide (1.0 equiv, 4.50 mmol, 517.6 mg) was added to an oven-dried 25 mL flask, which was evacuated and flushed with $N_2$ trice, followed by the addition of 5.0 mL THF. After cooling down in ice bath, 2-azidoacetic acid (1.0 equiv, 4.50 mmol, 336.7 μL) was added and stirred for 10 min at 0 °C. DCC (1.0 equiv, 4.50 mmol, 928.5 mg) was suspended in 5.0 mL THF and added dropwise. The reaction mixture was stirred for 4 h at 0 °C, followed by filtration to remove formed *N*,*N*'-dicyclohexylurea. 50 mL $Et_2O$ was added to the filtrate, which was stored at 4 °C overnight. The forming white solid was collected, rinsed with $Et_2O$ (20 mL), and dried to give 669.0 mg of **9** as white crystal solid in 75% yield. **$^1$H NMR** (400 MHz, $CDCl_3$) δ 4.24 (s, 2H, H-2), 2.87 (s, 4H, H-3', H-4'). **$^{13}$C NMR** (100 MHz, $CDCl_3$) δ 168.5 (C-2', C-5'), 164.3 (C-1), 48.1 (C-2), 25.7 (C-3', C-4'). **HR-MS** *m*/*z* calcd for $C_6H_7N_4O_4^+$ $(M + H)^+$ 199.0467, found 199.0460, deviation 3.5 ppm.

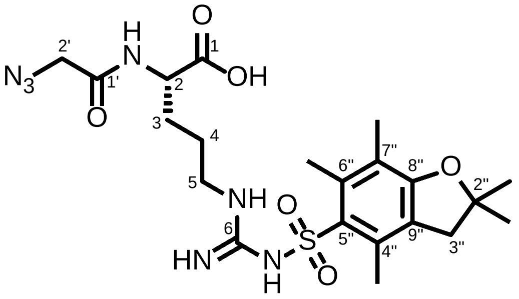

$N^2$-(2-Azidoacetyl)-$N^ω$-((2,2,4,6,7-pentamethyl-2,3-dihydrobenzofuran-5-yl)sulfonyl)-L-arginine **10**

In an oven-dried 50 mL flask, Fmoc-Arg(Pbf)-OH (1.0 equiv, 2.00 mmol, 1.298 g) was suspended in 30 mL DCM, followed by the addition of piperidine (5.0 equiv, 10.0 mmol, 987.8 μL). The reaction mixture was stirred for 16 h at room temperature and concentrated under reduced pressure. The resulting residue was added MTBE to yield white precipitate, which was collected by filtration and washed with MTBE to give Fmoc deprotected intermediate Arg(Pbf)-OH. The white solid was dried under vacuum and used without further purification. DMAP (0.1 equiv, 0.20 mmol, 24.4 mg) and compound **52** (1.5 equiv, 3.00 mmol, 594.1 mg) was added to the flask containing Arg(Pbf)-OH, which was evacuated and flushed with $N_2$ trice, followed by the addition of 10 mL DMF and $Et_3N$ (2.5 equiv, 5.00 mmol, 695.0 μL). After stirring overnight at room temperature, DMF was removed, and the residue was pre-purified via silica chromatography (DCM:MeOH = 20:1 ~ 3:1). Further RP-HPLC purification gave 472.2 mg **10** as pale yellow solid in 46% yield over two steps. **$^1$H NMR** (400 MHz, DMSO-$d_6$) δ 12.76 (br s, 1H, OH-1), 8.36 (d, *J* = 7.6 Hz, 1H, NH-2), 6.73-6.42 (m, 3H, -N<u>*H*</u>C(N<u>*H*</u>)N<u>*H*</u>-), 4.21-4.16 (m, 1H, H-2), 3.85 (s, 2H, H-2'), 3.04 (q, *J* = 6.5 Hz, 2H, H-5), 2.96 (s, 2H, H-3"), 2.48 (s, 3H, $CH_3$-6"), 2.42 (s, 3H, $CH_3$-4"), 2.01 (s, 3H, $CH_3$-7"), 1.76-1.67 (m, 1H, $H_a$-3), 1.60-1.51 (m, 1H, $H_b$-3), 1.45-1.41 (m, 2H, H-4), 1.41 (s, 6H, $CH_3$-2"). **$^{13}$C NMR** (100 MHz, DMSO-$d_6$) δ 173.0 (C-1), 167.4 (C-1'), 157.4 (C-8"), 156.1 (C-6), 137.2 (C-7"), 134.2 (C-5"), 131.4 (C-4"), 124.3 (C-9"), 116.3 (C-6"), 86.3 (C-2"), 51.8 (C-2), 50.4 (C-2'), 42.5 (C-3"), 39.6 (C-5, overlapped by DMSO-$d_6$), 28.4 (C-3), 28.3 ($CH_3$-2"), 25.6 (C-4), 18.9 ($CH_3$-4"), 17.6 ($CH_3$-6"), 12.3 ($CH_3$-7"). **HR-MS** *m*/*z* calcd for $C_{21}H_{30}N_7O_6S^-$ $(M - H)^-$ 508.1978, found 508.1985, deviation 1.4 ppm.

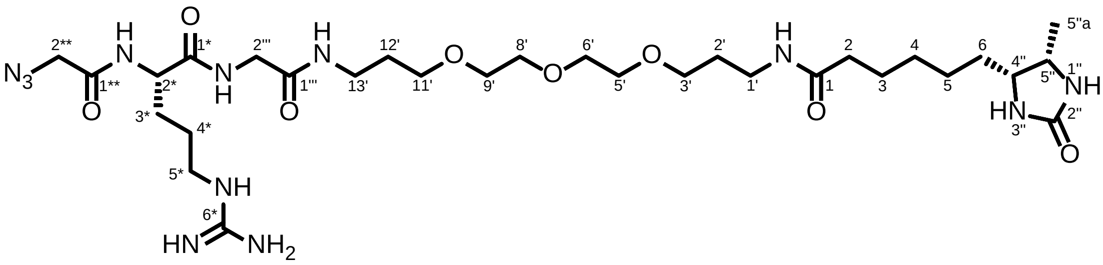

*N*-((*S*)-1-Amino-6-(2-azidoacetamido)-1-imino-7,10-dioxo-15,18,21-trioxa-2,8,11-triazatetracosan-24-yl)-6-((4*R*,5*S*)-5-methyl-2-oxoimidazolidin-4-yl)hexanamide

**$N_3$-(Arg)PEG$_3$-DB**

An oven-dried 10 mL flask was evacuated and flushed with $N_2$ trice, followed by the addition of compound **8** (1.0 equiv, 0.30 mmol, 1.10 mL 0.273 M solution in DMF) and $Et_3N$ (1.0 equiv, 0.30 mmol, 41.7 μL). Meanwhile, compound **9** (2.0 equiv, 0.60 mmol, 305.5 mg) and HOBt·x$H_2O$ (2.0 equiv, 0.60 mmol, 91.8 mg) was added to another oven-dried 10 mL flask, which was evacuated and flushed with $N_2$ trice, followed by the addition of 3.0 mL DMF. Then DCC (2.0 equiv, 0.60 mmol, 92.9 μL) was added and stirred for 15 min at room temperature, which was transferred to the flask containing compound **8** and $Et_3N$. After stirring for 3 h at room temperature, DMF was removed, and the residue was roughly purified via RP-HPLC. All fractions that contained Pbf-protected product were pooled, concentrated, and lyophilized. The crude product was then dissolved in 1 M HCl in HFIP and stirred for 8 h at room temperature to remove Pbf group. The reaction mixture was concentrated and purified via RP-HPLC with C18 Pyramid column using 0.1% TFA MilliQ® $H_2O$/0.1% TFA MeCN system. All fractions that contained Pbf-deprotected product were pooled, concentrated, and further purified via RP-HPLC with C18 column using MilliQ® $H_2O$ /MeCN system to remove excess amount of TFA to give 126.6 mg of **$N_3$-(Arg)PEG$_3$-DTB** as colorless oil in 59% yield over two steps. **$^1$H NMR** (400 MHz, DMSO-$d_6$) δ 8.52 (d, *J* = 7.2 Hz, 1H, NH-2$^*$), 8.40 (t, *J* = 5.8 Hz, 1H, NH-2'''), 7.82-7.75 (m, 3H, NH-1', NH-13', NH-3''), 7.44-7.19 (m, 5H, NH-1'', -N*H*C(N*H*)N*H*$_2$), 4.25 (q, *J* = 6.8 Hz, 1H, H-2$^*$), 3.90 (s, 2H, H-2$^{**}$), 3.72-3.58 (m, 3H, H-5'', H-2'''), 3.51-3.45 (m, 9H, H-5', H-6', H-8', H-9', H-4''), 3.40-3.36 (m, 4H, H-3', H-11'), 3.12-3.03 (m, 6H, H-1', H-13', H-5$^*$), 2.04 (t, *J* = 7.4 Hz, 2H, H-2), 1.77-1.15 (m, 16 H, H-3, H-4, H-5, H-6, H-2', H-12', H-3$^*$, H-4$^*$), 0.96 (d, *J* = 6.0 Hz, 3H, H-5''a). **$^{13}$C NMR** (100 MHz, DMSO-$d_6$) δ 172.0 (C-1), 171.3 (C-1$^*$), 168.5 (C-1'''), 167.9 (C-1$^{**}$), 162.8 (C-2''), 157.0 (C-6$^*$), 69.8 (C-6', C-8'), 69.5 (C-5', C-9'), 68.1 (C-3'), 68.0 (C-11'), 55.0 (C-4''), 52.7 (C-2$^*$), 50.5 (C-2$^{**}$), 50.3 (C-5''), 42.1 (C-2'''), 40.3 (C-5$^*$), 35.9 (C-13'), 35.7 (C-1'), 35.3 (C-2), 29.5 (C-6), 29.4 (C-2'), 29.2 (C-12'), 28.8 (C-4), 28.7 (C-3$^*$), 25.6 (C-5), 25.2 (C-3), 25.0 (C-4$^*$), 15.5 (C-5''a). **HR-MS** *m*/*z* calcd for $C_{30}H_{57}N_{12}O_8^+$ $(M + H)^+$ 713.4422, found 713.4422, deviation 0.0 ppm.

#### NMR Spectra

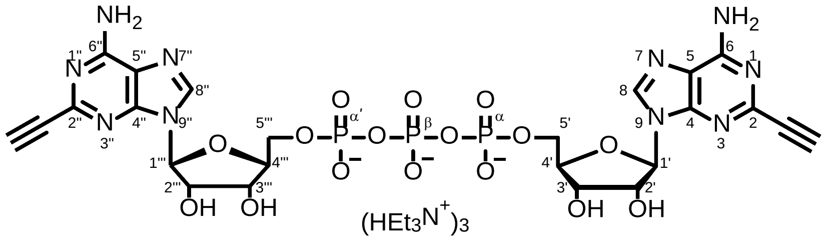

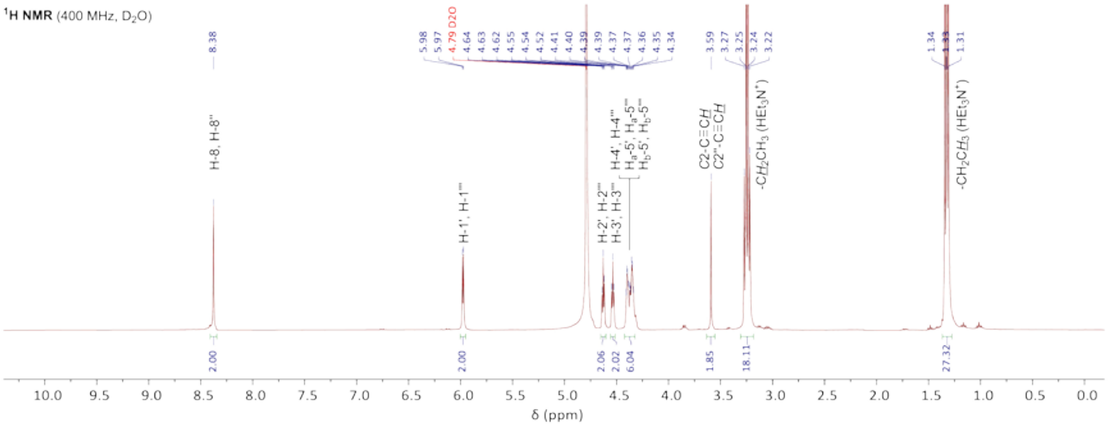

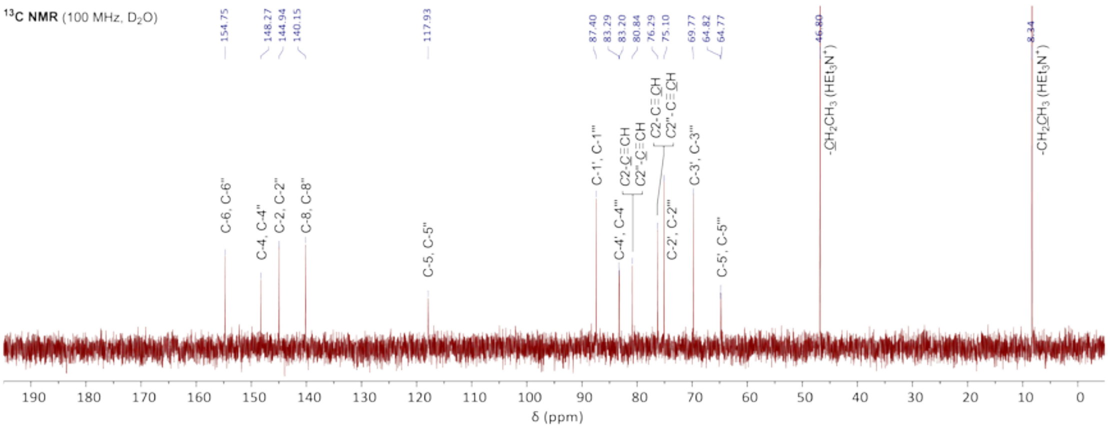

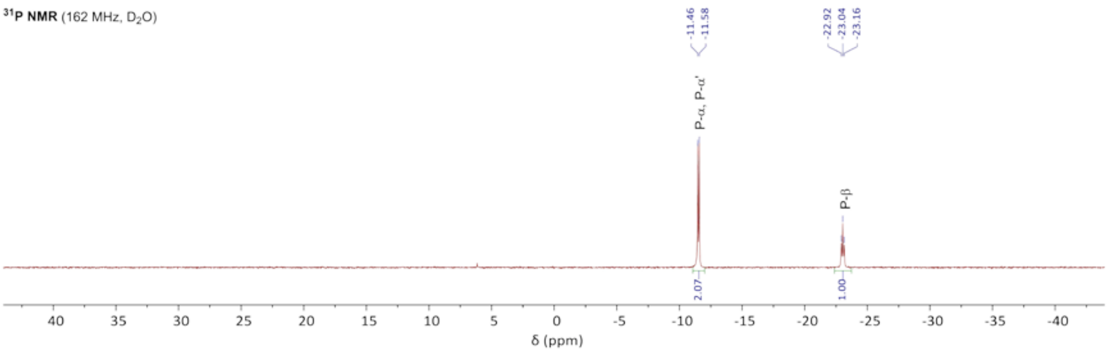

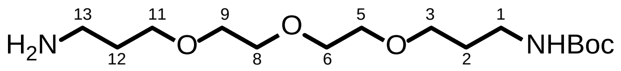
H2N
NHBoc
O
O
O
13
12
11
9
8
5
6
3
2
1

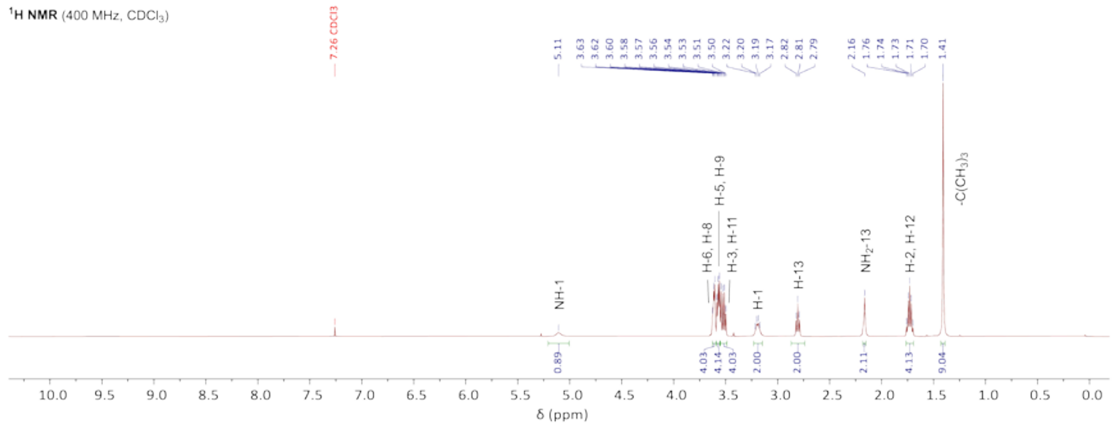
1H NMR (400 MHz, CDCl3)
7.26 CDCl3
NH-1
H-6, H-8
H-5, H-9
H-3, H-11
H-1
H-13
NH2-13
H-2, H-12
-C(CH3)3
δ (ppm)

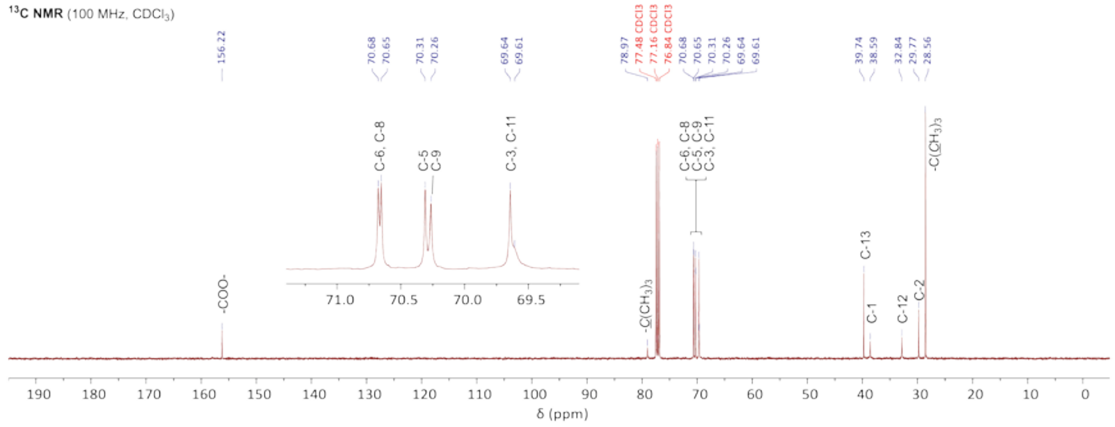
13C NMR (100 MHz, CDCl3)
156.22
77.48 CDCl3
77.16 CDCl3
76.84 CDCl3
-COO-
-C(CH3)3
C-6, C-8
C-5, C-9
C-3, C-11
C-13
C-1
C-12
C-2
-C(CH3)3
δ (ppm)

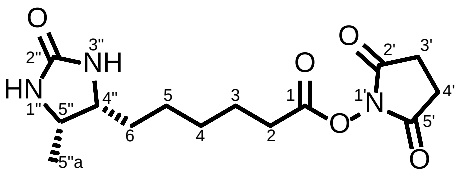

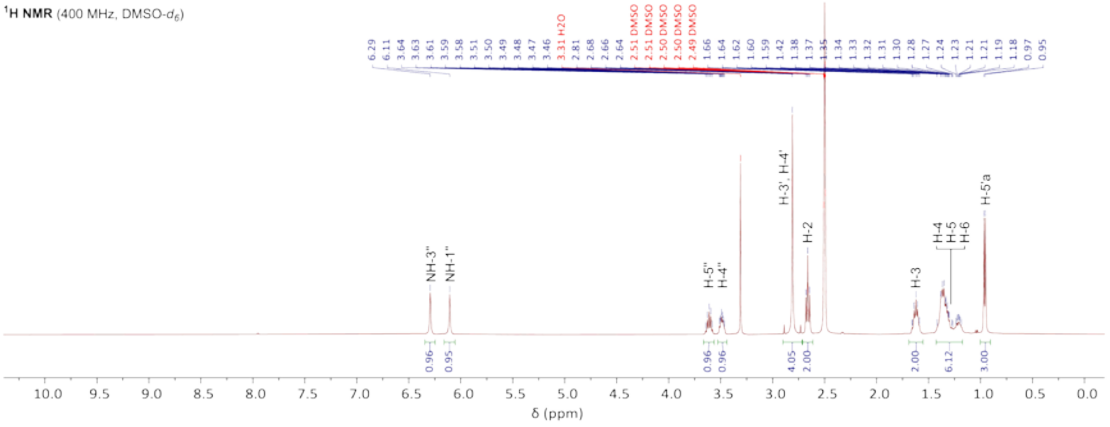

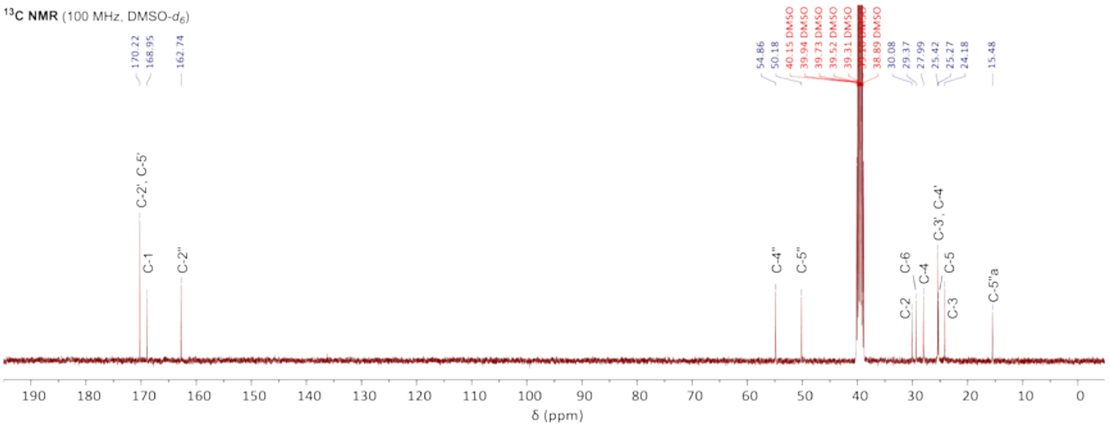

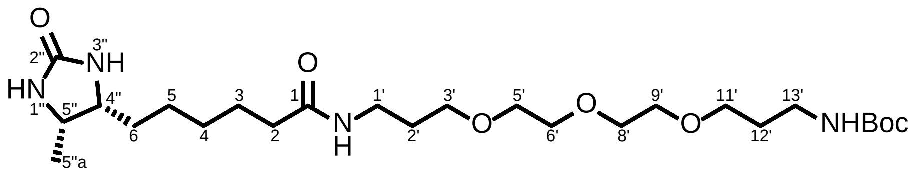

$^{1}$**H NMR** (400 MHz, DMSO-$d_6$)

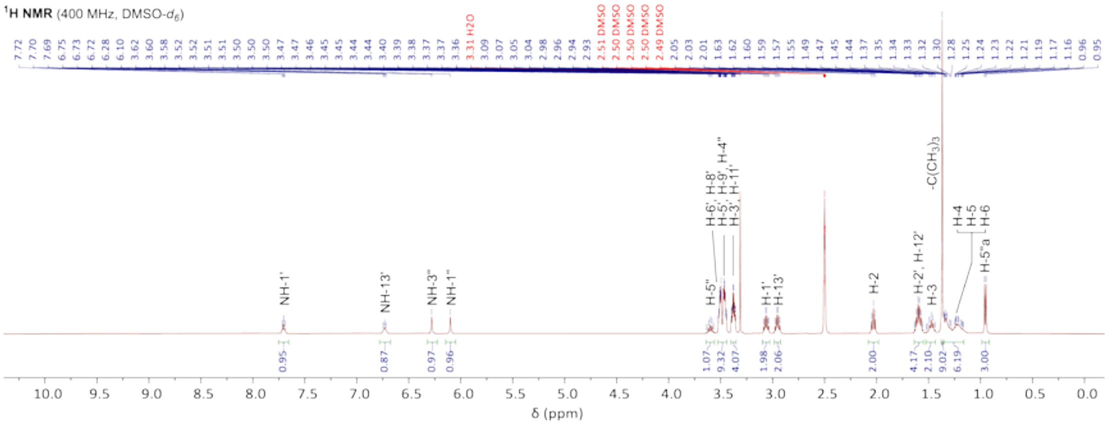

$^{13}$**C NMR** (100 MHz, DMSO-$d_6$)

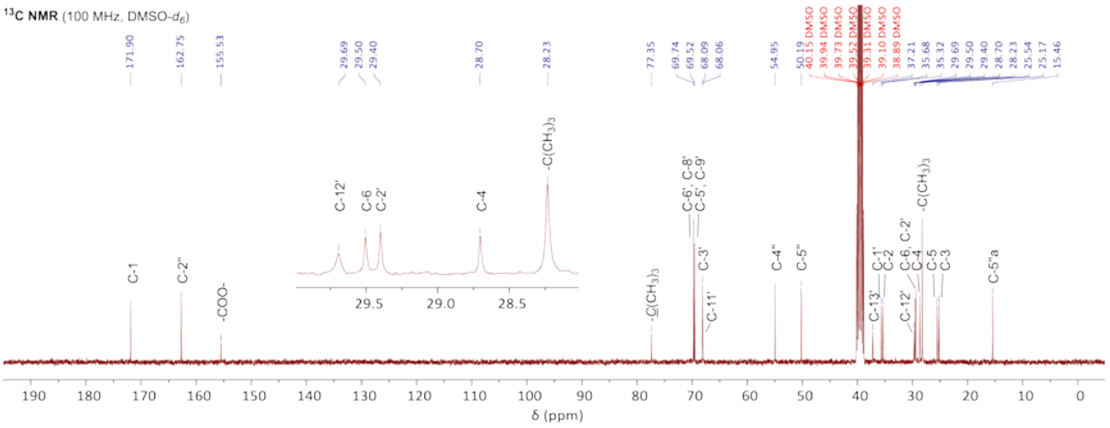

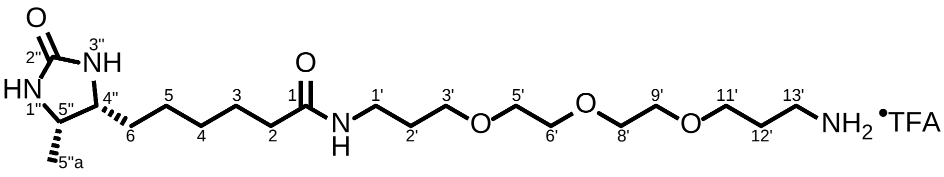

$^{1}$H NMR (400 MHz, DMSO-$d_6$)

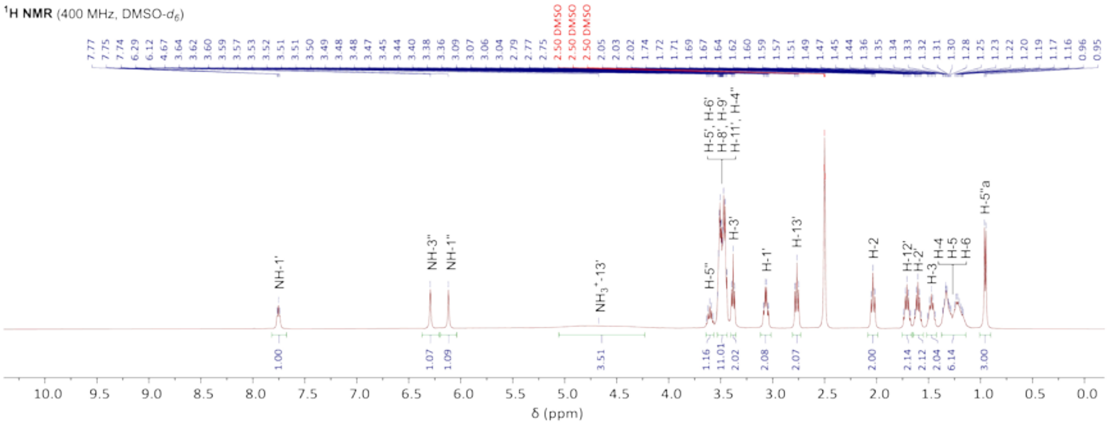

$^{13}$C NMR (100 MHz, DMSO-$d_6$)

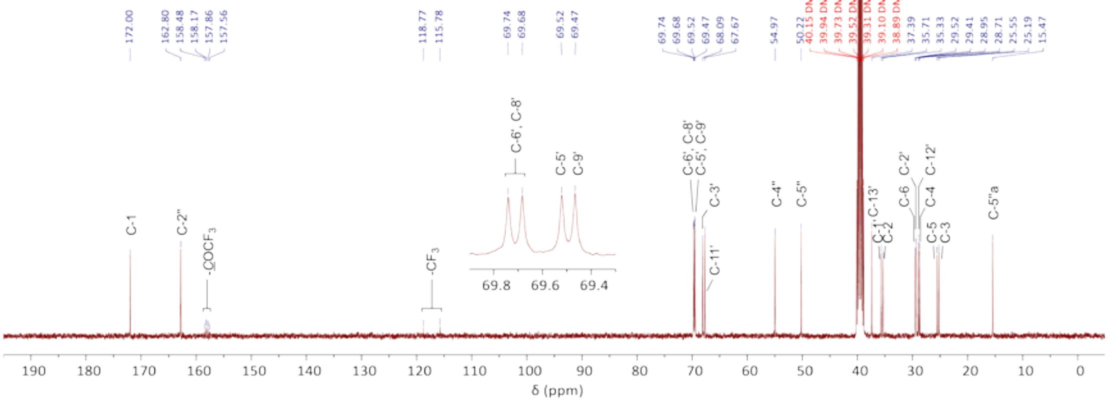

$^{19}$F NMR (376 MHz, DMSO-$d_6$)

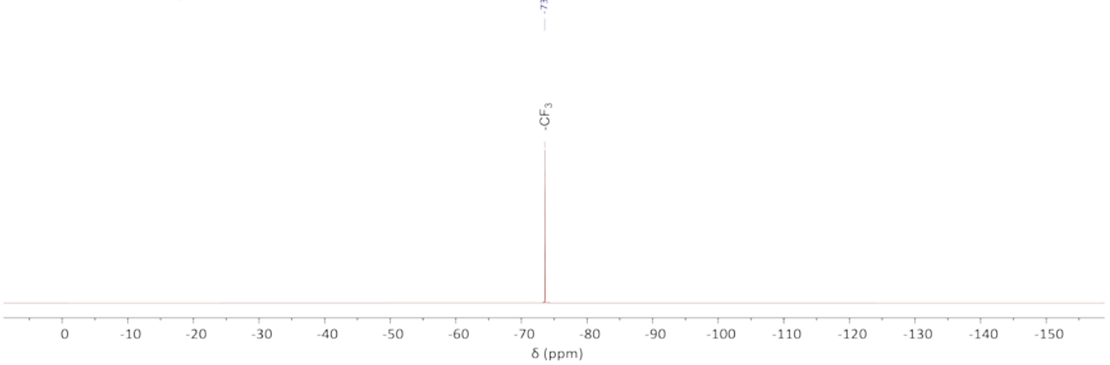

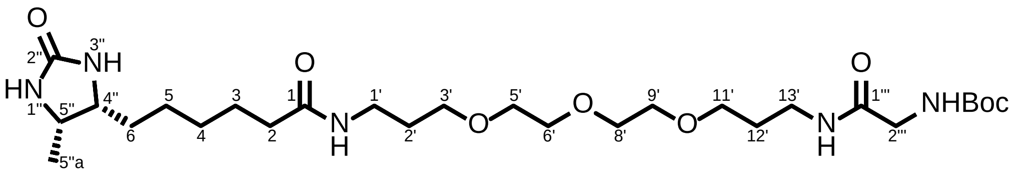

$^{1}$H NMR (400 MHz, DMSO-$d_6$)

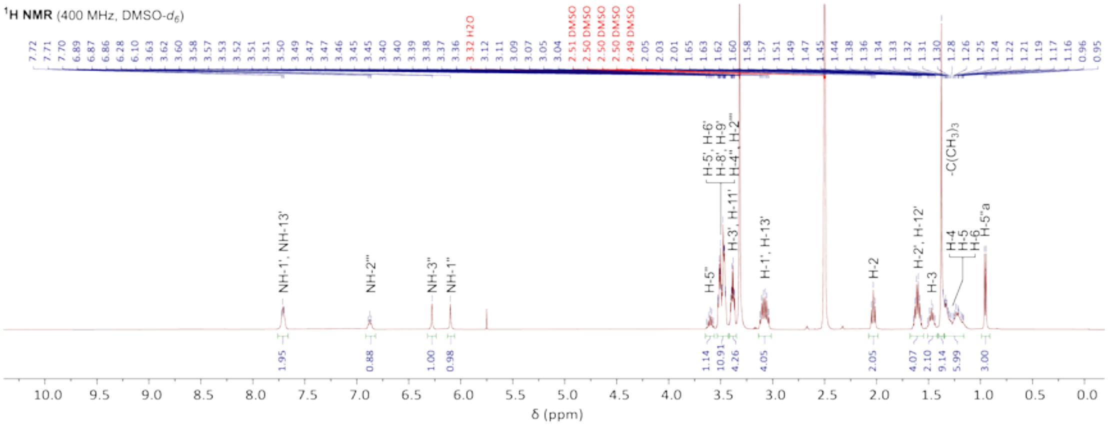

$^{13}$C NMR (100 MHz, DMSO-$d_6$)

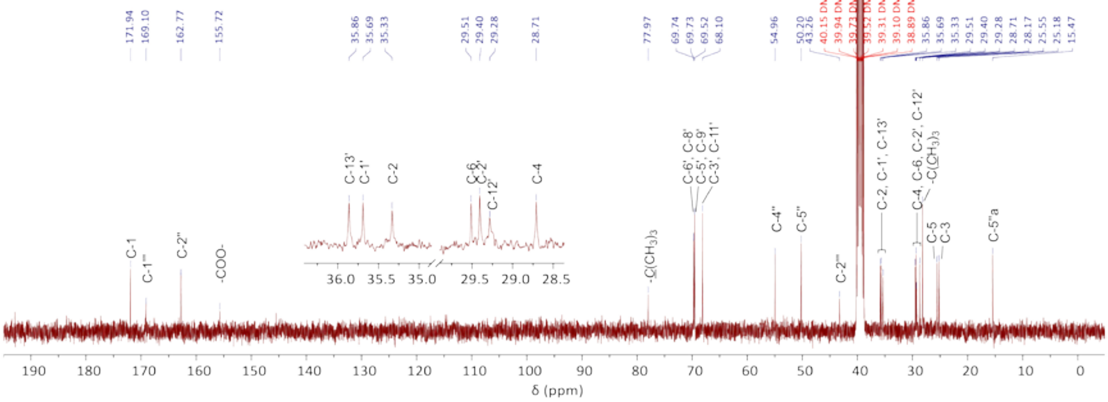

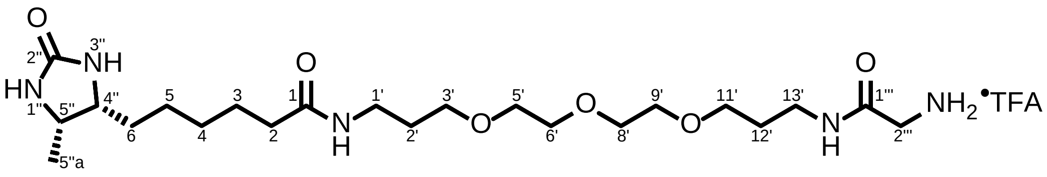

$^1$H NMR (400 MHz, DMSO-$d_6$)

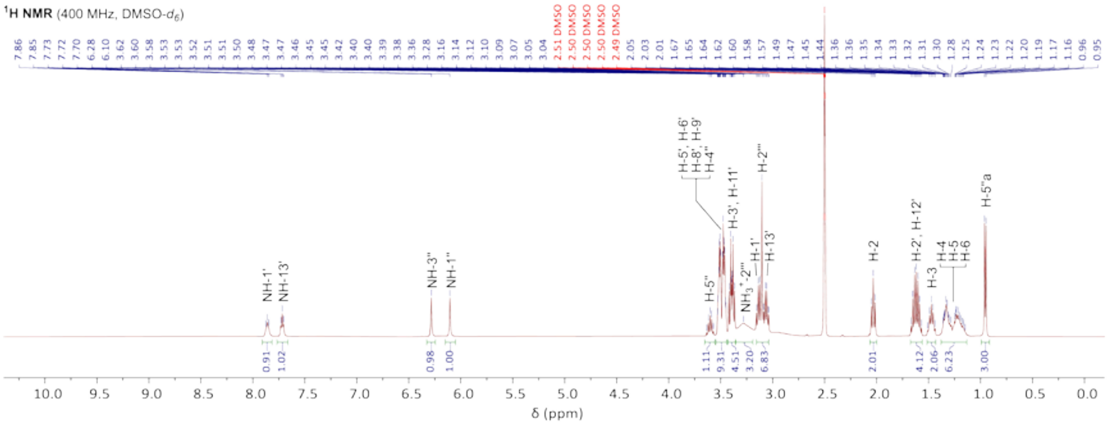

$^{13}$C NMR (100 MHz, DMSO-$d_6$)

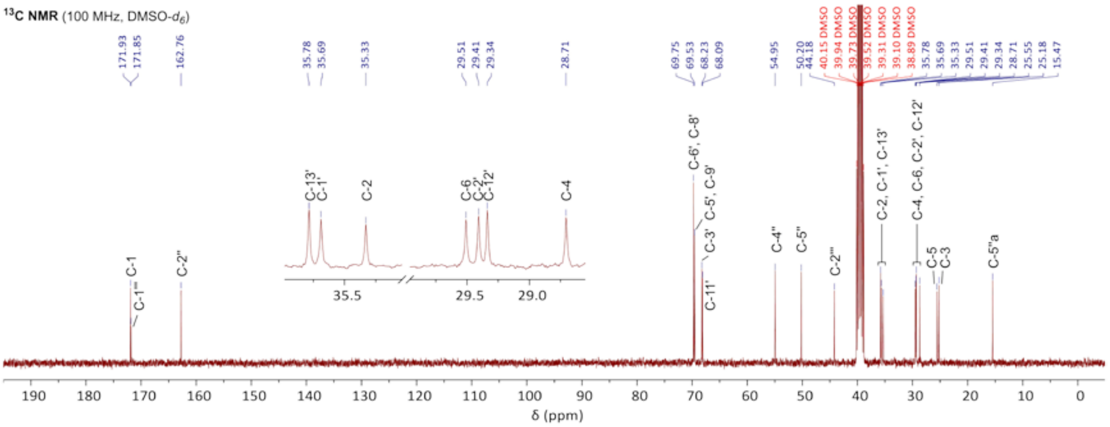

$^{19}$F NMR (376 MHz, DMSO-$d_6$)

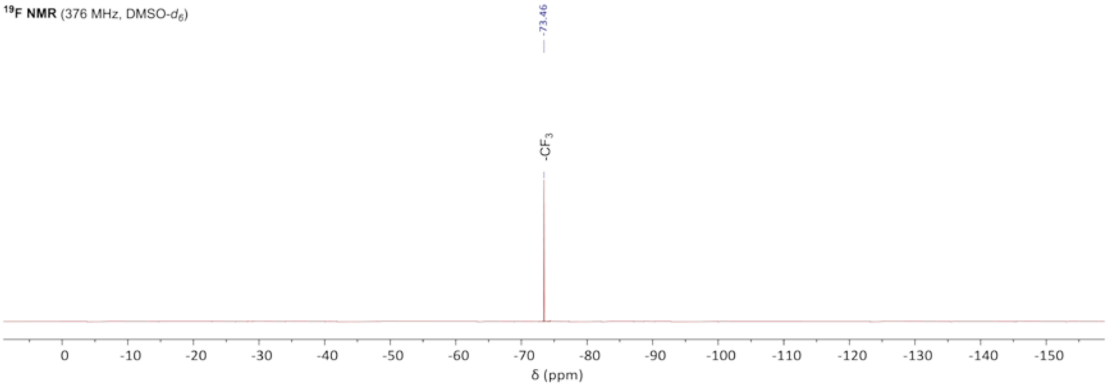

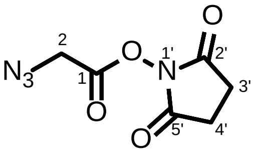

$^{1}$**H NMR** (400 MHz, $CDCl_3$)

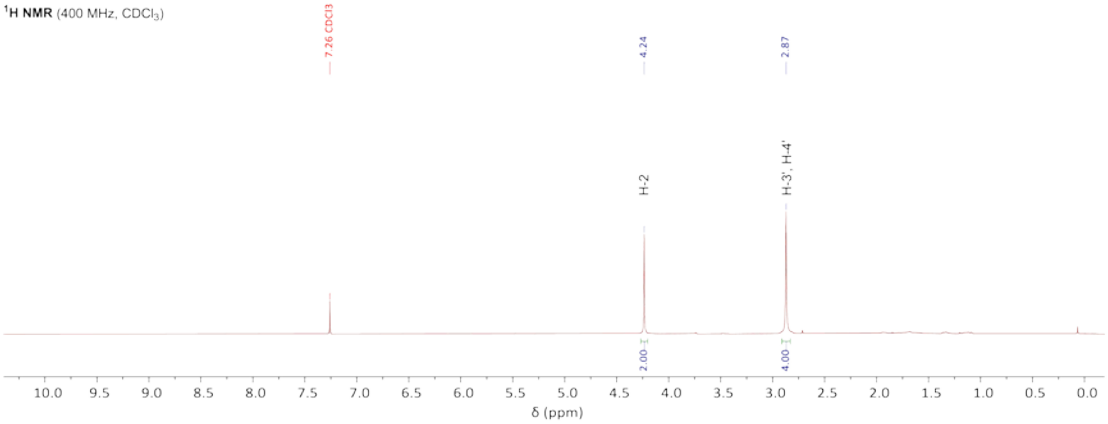

$^{13}$**C NMR** (100 MHz, $CDCl_3$)

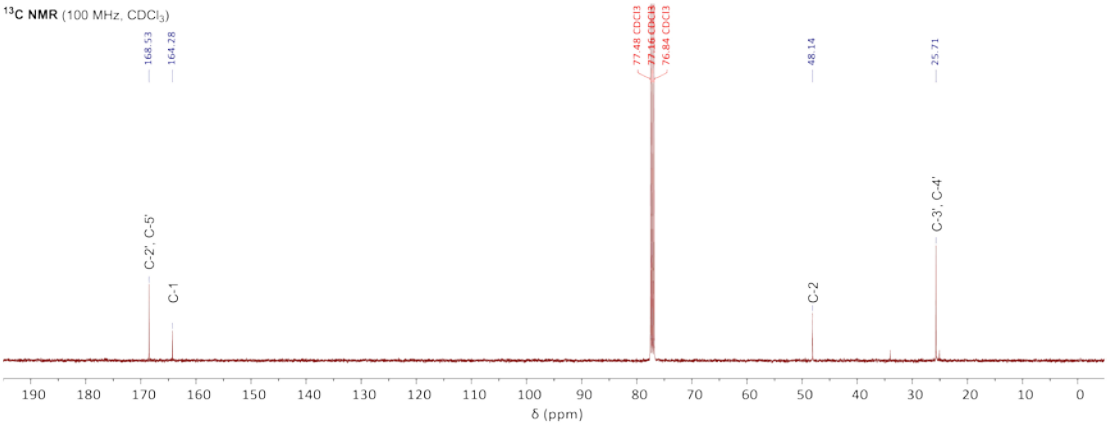

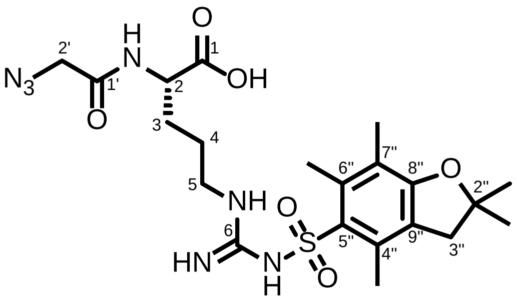
N3
H
N
O
1
OH
2'
1'
2
O
3
4
5
NH
6
HN
N
H
S
O
O
6"
7"
8"
O
2"
5"
4"
9"
3"

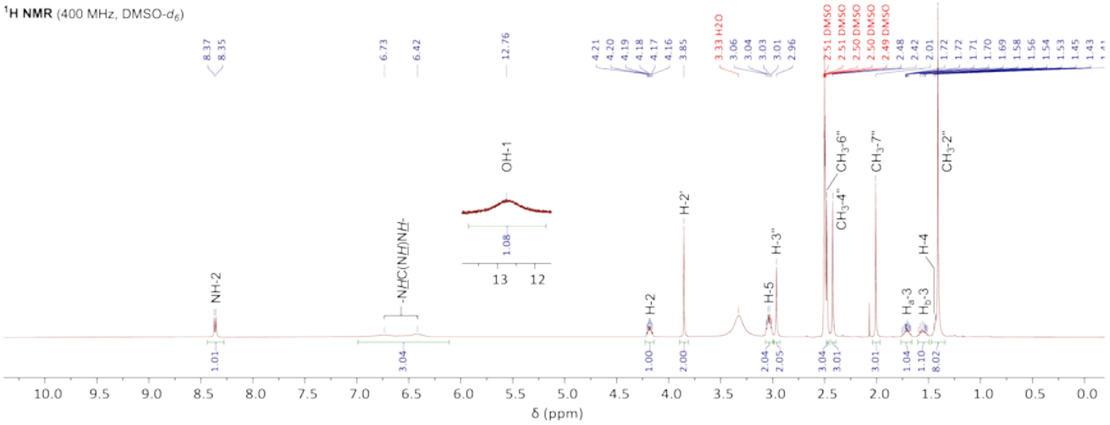
1H NMR (400 MHz, DMSO-d6)
NH-2
-NHC(NH)NH-
OH-1
H-2
H-2'
H-5
H-3"
CH3-6"
CH3-4"
CH3-7"
Ha-3
Hb-3
H-4
CH3-2"
δ (ppm)

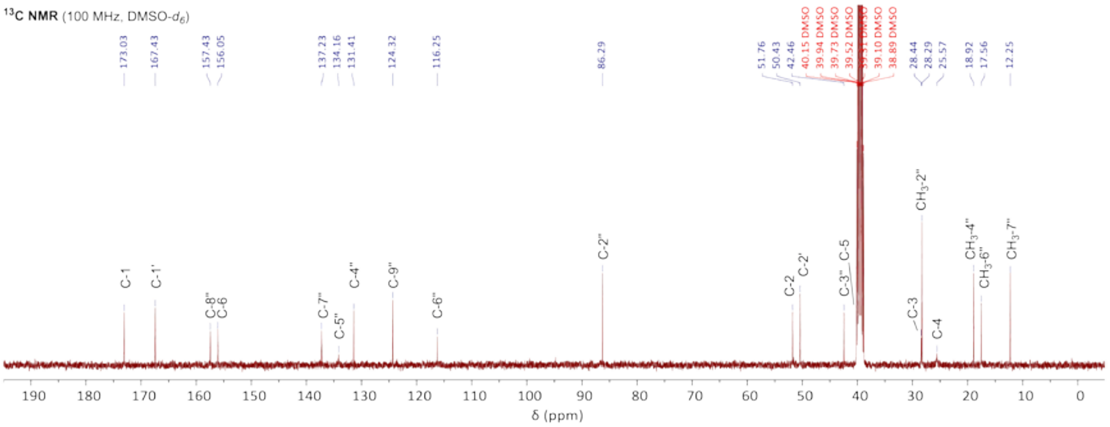
13C NMR (100 MHz, DMSO-d6)
C-1
C-1'
C-8"
C-6
C-7"
C-5"
C-4"
C-9"
C-6"
C-2"
C-2
C-2'
C-3"
C-5
C-3
C-4
CH3-2"
CH3-4"
CH3-6"
CH3-7"
δ (ppm)

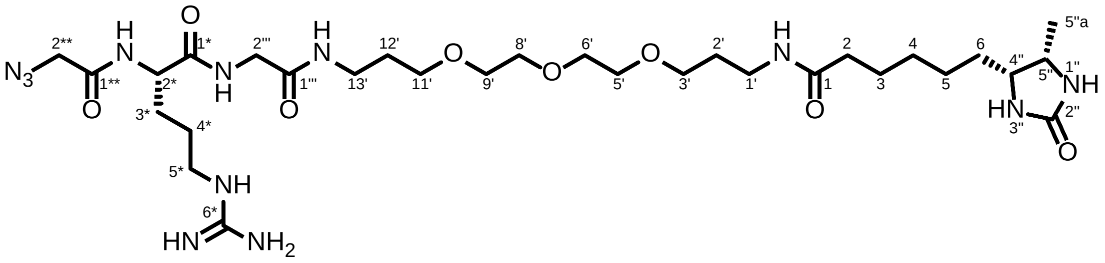

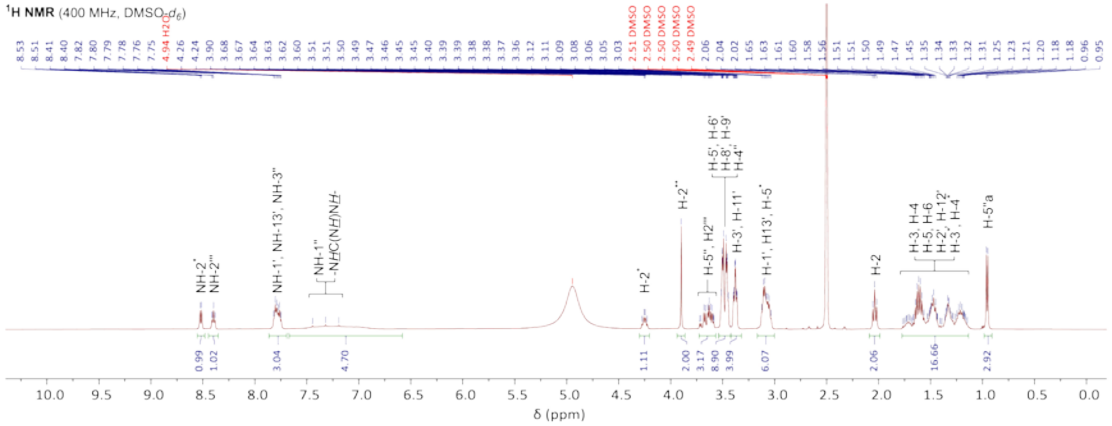
1H NMR (400 MHz, DMSO-d6)
δ (ppm)

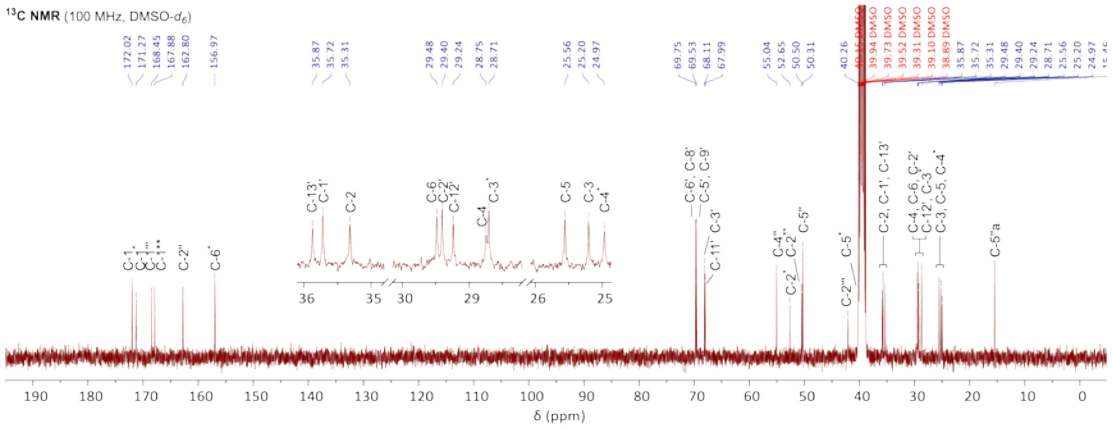
13C NMR (100 MHz, DMSO-d6)
δ (ppm)
